# Modularity and evolution of flower shape: the role of efficiency, development, and spandrels in *Erica*

**DOI:** 10.1101/628644

**Authors:** Dieter Reich, Andreas Berger, Maria von Balthazar, Marion Chartier, Mahboubeh Sherafati, Christian P. Klingenberg, Sara Manafzadeh, Yannick M. Staedler

**Author notes:** Equal contribution (Dieter Reich, Andreas Berger). Corresponding author: Yannick M. Staedler.

## Abstract

- Three hypotheses can explain floral modularity: the attraction-reproduction, the efficiency, and the developmental hypotheses.
- In order to test these hypotheses and understand if pollination specialisation and pollination syndrome influence floral modularity, we focussed on the genus *Erica*: we gathered 3D data from flowers of species with diverse pollination syndromes via Computed Tomography, and analysed their shape via geometric morphometrics. In order to provide an evolutionary framework for our results we tested the evolutionary mode of floral shape, size, and integration under pollination syndrome regimes, and - for the first time-reconstructed the high-dimensional floral shape of their most recent common ancestor.
- We demonstrate, for the first time, that the modularity of generalist flowers depends on development and that of specialists is linked to efficiency: in bird syndrome flower, efficiency modules were associated with pollen deposition and receipt, whereas in long-proboscid fly syndrome, they were associated with restricting the access to the floral reward. Only shape PC1 showed selection towards multiple optima, suggesting that PC1 was co-opted by evolution to adapt flowers to novel pollinators. Whole floral shape followed an OU model of evolution, and demonstrated relatively late differentiation.
- Flower shape modularity thus crucially depends on pollinator specialisation and class.

## Introduction

From the bacterial flagellum (McAdams *et al.*, 2004) to the skull shape of dinosaurs (Fabbri *et al.*, 2017), modular organisation pervades life’s phenotype (Wagner *et al.*, 2007). Modules are subsets of traits that are integrated (i.e. they tend to vary in a coordinated manner) that vary relatively independently from other such subsets (Klingenberg, 2014). Relative independence of modules allows for evolutionary tinkering to take place in one module without much affecting the other (Alon, 2003; Kirsten & Hogeweg, 2011). Modular organisation is thus not only a key feature of the structural complexity of life, but also a key feature for its evolvability (Wagner *et al.*, 2007). Theophrastus’ observation, twenty three centuries ago, that “repetition is of the essence of plants” (Theophrastus & Hort, 1916) is underlain by plants’ non-conformity to Weissman’s doctrine of separation of soma and germ (Weismann, 1892): the indefinite developmental totipotency of meristematic plant cells allows for the modular construction of plants by continuous organogenesis and the repeated production of homologous structures (Herrera, 2009). However, despite the fundamentally modular structure of plants (see Ottaviani et al. 2017 and references therein), historically, most studies of modularity have, and still are, focussed on animals (Klingenberg, 2014; Esteve-Altava, 2017)(see Notes S1). In her seminal work, Raissa Berg hypothesised that the variation of traits in specialised flowers is largely uncorrelated with that of vegetative traits (Berg, 1960), i.e. that vegetative and reproductive traits form independent modules, which are themselves highly integrated (Wagner & Altenberg, 1996). Because different floral traits can experience different selection pressures, Berg’s hypothesis can be expanded to include modules of traits within the flower (Ordano *et al.*, 2008; Diggle, 2014; Armbruster & Wege, 2018).

Accordingly, the following explicit hypotheses of flower modularity have been advanced. The first hypothesis is the *attraction-reproduction modularity hypothesis*; this hypothesis proposes that flowers are divided into a module of attraction comprising the petals and the sepals, and a module of reproduction comprising the stamens and the carpels (Esteve-Altava, 2017), see Fig. 1a. The second general hypothesis is the *efficiency modularity hypothesis*, which proposes that flowers are divided into a module efficiency that comprises parts from different organs that effect reproduction (constriction of floral tube, pollen sacs of the stamens, stigma of the carpels, etc.), and a module of attraction (e.g. showy part of petals)(Diggle, 2014). This hypothesis has been supported by multiple studies (Herrera, 2001; Fenster *et al.*, 2004; Pigliucci & Preston, 2004; Carvallo & Medel, 2005; Pérez *et al.*, 2007; Bissell & Diggle, 2008; Ordano *et al.*, 2008; Bissell & Diggle, 2010; Fornoni *et al.*, 2015; Heywood *et al.*, 2017; Armbruster & Wege, 2018). *Efficiency hypotheses* can comprise modules of pollen deposition and receipt (see efficiency 1 in Fig. 1b), or modules involving putative pollinator filters such as corolla aperture (see efficiency 2 in Fig. 1c).

**Figure 1.**
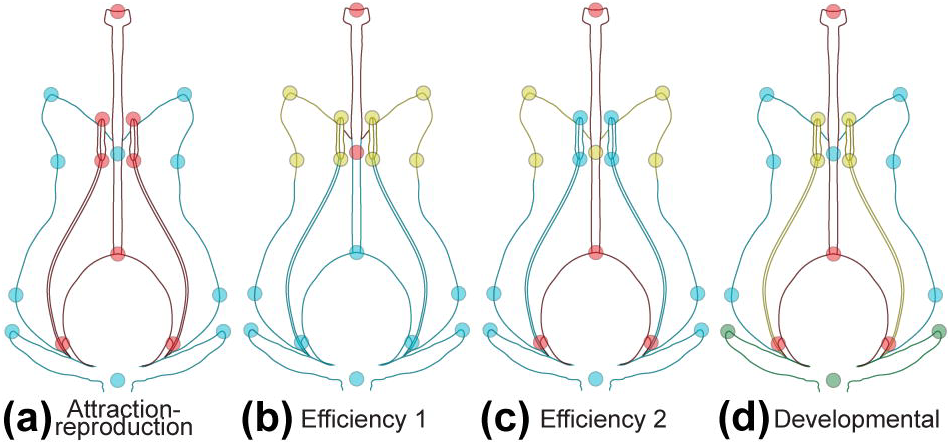
Hypotheses. Modularity hypotheses tested displayed on schematic representation of an *Erica* flower. (a) the *attraction-reproduction* hypothesis proposes that floral organs groups into fertile (stamens and carpel, in red) versus sterile (sepals and petals, in blue) modules. (b) the *efficiency* hypothesis 1 proposes that parts of the flower group in modules directly involved in pollen receipt (joining of the petals and stigma, in red) and deposition (rest of the corolla mouth and stamens, in yellow), and modules that are not (remainder of the flower in blue). (c) the *efficiency* hypothesis 2 proposes that parts of the flower that restrict access to the floral reward (floral neck, in yellow) form a module, that the carpels form a module, and that the rest of the flower also forms a module. (d) the *developmental* hypothesis proposes that parts for the flower group into modules corresponding to their organ identity: sepals (green), petals (blue), stamens (yellow), or carpels (red).

The abovementioned two hypotheses fail to incorporate the specificities of flowers and their fundamental difference from animal structures. Modules in animals are typically searched for on different parts of the same organ, such as the skull (e.g. (Drake & Klingenberg, 2010; Bardua *et al.*, 2019)), the jaw (e.g. (Hulsey *et al.*, 2006)), or the wing (e.g. (Klingenberg *et al.*, 2010; Chazot *et al.*, 2016)), whereas flowers are complexes of fundamentally different organs performing fundamentally different functions, such as protection from predators (i.e. sepals), sexual attraction (i.e. petals), male reproduction (i.e. stamens), and female reproduction (i.e. carpels). Moreover, despite their current functional association, these organs have (mostly) evolved from different progenitors and most likely without functional association for ca. 125 million years, from the origin of seed plants to that of flowering plants (Morris *et al.*, 2018). We thus propose a third explicit hypothesis of modularity in flowers: *the developmental modularity hypothesis* proposes that floral modularity is dominated by developmental factors, i.e. that each organ class (sepal, petal, stamen, and carpel) forms its own module (see Fig. 1d). The converse of floral modularity, the tendency of groups of features to be independent from each other, is whole-flower integration, the tendency for all the features of the flower to covary. Whole-flower integration level has been hypothesised to vary according to pollination system (see below).

Flowers are pollinated by organisms that differ greatly in their morphology and sensory systems (see e.g. (Kelber & Jacobs, 2016)). This has led to convergences in the floral morphology of species pollinated by the same group(s) of animals, as described in the pollination syndrome hypothesis (Vogel, 1954; Grant & Grant, 1965; Stebbins, 1970; Johnson, 2006). Syndromes can be divided into specialised syndromes (pollination by one group of pollinators), and generalised syndromes (pollination by several groups of pollinators). In flowers with specialised syndromes, we expect to observe support for different versions of the *efficiency modularity* (see, e.g. (Diggle, 2014)), depending on pollinator class. In generalist flowers, however three main hypotheses have been advanced to explain how pollinators affect floral shape (Aigner, 2001; Sahli & Conner, 2011; Joly *et al.*, 2018), each of which would lead to different floral modules. (1) The “trade-off” hypothesis (Aigner, 2001; Aigner, 2006; Sahli & Conner, 2011) suggests that a change in trait that increases the fitness contribution of one pollinator will decrease the fitness of another. This model predicts that selection by multiple pollinators in multiple directions would cancel each other out, resulting in weak or absent *efficiency modularity*, in which case *developmental modularity* should be observed instead. (2) The “trait specialisation” hypothesis (Sahli & Conner, 2011) proposes that individual traits are under selection by a subset of pollinators, resulting in flowers that possess different traits adapted to different pollinators (which predicts several, well-defined *efficiency modules*). (3) The “common shape” hypothesis (Sahli & Conner, 2011) implies that the different pollinators all select for a common shape, which also predicts the existence of *efficiency modules*.

Specialisation and pollinator groups have been hypothesised and shown to also influence whole-flower integration (hereafter only referred to as “integration”). That flowers are highly integrated organ complexes has become a paradigm among floral biologists (Stebbins, 1950; Faegri & Van Der Pijl, 1966; Stebbins, 1970; Ordano *et al.*, 2008), as is the hypothesis that specialised flowers are more highly integrated than generalist flowers because specialised pollination is expected to drive the evolution of precise, highly coordinated (integrated) floral traits (Armbruster *et al.*, 1999; Pérez *et al.*, 2007; Rosas-Guerrero *et al.*, 2011; Ellis *et al.*, 2014; Gomez *et al.*, 2014; Gomez *et al.*, 2016). Support for this hypothesis has been provided (Meng *et al.*, 2008; Rosas-Guerrero *et al.*, 2011; Gomez *et al.*, 2014); however, support for the opposite hypothesis has unexpectedly also been provided (Armbruster *et al.*, 1999; Edwards & Weinig, 2011; Joly *et al.*, 2018). Moreover, the group of pollinators possibly also determines the magnitude of integration of the flowers (Pérez-Barrales *et al.*, 2007; Gomez *et al.*, 2014; Pérez-Barrales *et al.*, 2014; González *et al.*, 2015).

Natural selection is an optimising mechanism that increases the accuracy of complex traits, increasing their precision and decreasing their variation (Bell, 1997; Hansen *et al.*, 2006; Gomez *et al.*, 2016). In specialist flowers, floral shape and size should show evidence of stabilising selection around an optimal shape and size adapted to its pollinator, whereas in generalists, the trade-off hypothesis predicts relaxed selection constraints (Johnson & Steiner, 2000), and both the trait specialisation hypothesis and the common shape hypothesis (Sahli & Conner, 2011) predict selection similar to that present in specialists. Therefore, from a macro-evolutionary perspective, if floral shape, size, and integration are affected by pollination syndromes, we would expect that within a lineage where a number *N* of pollination syndromes evolved repeatedly, the evolution of these floral parameters follows a natural selection model such as an Orstein-Uhlenbeck (OU) process with *N* optima. Alternatively, if floral shape, size, and integration are not affected by pollination syndromes, we would expect that the evolution of these floral parameters follows a drift–like model such as the Brownian Motion (BM) process instead.

To our knowledge, no study has yet tested if floral modules change with pollinator groups; if modularity type changes with specialisation, and what the process of evolution of 3D shape, size, and integration in a system with convergent evolution of pollinator systems is. To answer these questions and test these hypotheses requires a study system in which convergent evolution of specialist pollination systems occurred, and that also contains species with generalist pollination; such a system should also possess a constant floral bauplan in order to rigorously homologise structures. *Erica* is such a system: it is a large genus of ca. 800 species mostly distributed in South Africa (Pirie *et al.*, 2016). Within the many South African members of the genus, evolution of pollination via birds and long-proboscid flies (LPF) has possibly repeatedly taken place (Pirie *et al.*, 2011), whereas generalist pollination syndrome has been found to be prevalent in European species (see Table 1). Moreover, the flowers of *Erica* have consistently the same, 4-merous bauplan with mostly 8 stamens (Stevens *et al.*, 2004). *Erica* is thus the ideal system to test the effects of pollinator shifts on floral modularity.

**Table 1.** Sampling, systematic syndrome, observed (a, b, e-n) or predicted (in the literature: c, d, or via machine learning: RF = Random Forests), and number of flowers scanned.

| **Species** | **Clade<sup>a</sup>** | **Syndrome** | **Reference** | **n (flowers)** |
| --- | --- | --- | --- | --- |
| *Erica australis* L. | Paelearctic | gen | b | 11 |
| *Erica blandfordia* Andrews | Cape | gen | c, RF | 11 |
| *Erica bolusiae* T. M. Salter | Cape | gen | c, RF | 10 |
| *Erica brachialis* Salisb. | Cape | bird | c, j, k | 14 |
| *Erica capensis* T.M. Salter | Cape | gen | c, n | 10 |
| *Erica curviflora* L. | Cape | bird | c, RF | 11 |
| *Erica georgica* L. Guthrie & Bolus | Cape | lpf | RF | 15 |
| *Erica gracilis* J.C. Wendl. | Cape | gen | l | 10 |
| *Erica hirtiflora* Curtis | Cape | gen | c, m | 10 |
| *Erica lateralalis* Willd. | Cape | gen | c, d, RF | 10 |
| *Erica leucotrachela* H.A. Baker | Cape | bird | c, RF | 10 |
| *Erica margaritacea* Aiton | Cape | gen | c, RF | 13 |
| *Erica melanthera* L. | Cape | gen | c, RF | 10 |
| *Erica perspicua* J.C. Wendl. | Cape | bird | e, c, g | 10 |
| *Erica scoparia* L. | Paelearctic | wind | f | 10 |
| *Erica spiculifolia* Salisb. | Paelearctic | gen | RF | 12 |
| *Erica turgida* Salisb. | Cape | gen | c, RF | 12 |
| *Erica vagans* L. | Paelearctic | gen | h, i | 11 |
| *Erica ventricosa* Thunb. | Cape | lpf | c, **d** | 9 |

Visitor data from literature, websites, and personal observation. gen: insect generalist pollination syndrome; LPF: long-proboscid fly. a, (Pirie *et al.*, 2016); b, (Gil-López *et al.*, 2014); c, (Rebello *et al.*, 1985); d, (Rebello *et al.*, 1984); e, (Heystek *et al.*, 2014); f, (Herrera, J, 1988); g, (Geerts, 2011); h, (Fern & Fern, 2012); i, (Plants\_Database, 2019); j, (Turner, 2010); k, (Notten, 2012); l, Yannick M. Staedler, pers. obs. on cultivated specimen; m, (Arendse, 2015); n, (Cullinan *et al.*). RF, syndrome predicted via random forests. c and d, contain description of syndromes and attribute different *Erica* species to them. \* contains mention of observation for this species.

In order to test the abovementioned modularity and macro-evolutionary hypotheses, we generated 3D models of *Erica* flowers, the shape of which we digitised using geometric morphometric landmarks. We then used this shape dataset to test our different modularity hypotheses in *Erica* flowers (attraction-reproduction, developmental, and efficiency 1 and 2) in flowers with different pollination syndromes. We used phylogenetic reconstructions to test if floral parameters (shape, size, and integration) evolved under selection driven by pollination syndromes or randomly. We thus aim to understand: (1) The relative importance of the components of floral shape and size in predicting pollination syndromes (2) How floral shape modularity changes with pollination syndromes and floral specialisation (3) The possible evolutionary patterns of floral shape in *Erica* (4) The relative roles of natural selection models (i.e. Ornstein-Uhlenbeck) and drift-like models (i.e. Brownian motion) in explaining the evolution of floral shape, size, and integration in respect to pollination syndromes.

## Materials and Methods

### Plant material

We analysed ca. 10 flowers each from a single genotype representing nineteen species of *Erica* from the collections of the greenhouses of the Belvedere Garden (Austrian Federal Gardens). We selected species based on their diversity in pollination syndrome (generalist, bird, long-proboscid flies, and wind) and broadly representative phylogenetic position. Although limited, our selection contains both older European lineages and species from the more recently diversified and species rich South-Western Cape Clade as defined by (Pirie *et al.*, 2011; Pirie *et al.*, 2016), see Method S1 and Table 1 for details.

### X-ray tomography

Flowers were contrasted, mounted, and scanned according to (Staedler *et al.*, 2013). See Methods S1 and Table S1 for details.

### 3D-landmarking & Geometric Morphometrics

Geometric morphometric landmarking was carried out on isosurface models in AMIRA. Thirty-three homologous landmarks were placed on each flower (see Fig. 2a-c, Table S2). Landmark coordinates were exported as csv-files, concatenated, and imported in MorphoJ 1.06d (Klingenberg, 2011). Procrustes fit, and calculation of the covariance matrix, Principal Component Analysis (PCA), modularity analyses, and allometric regressions were performed in MorphoJ. See Methods S1 for details.

**Figure 2.**
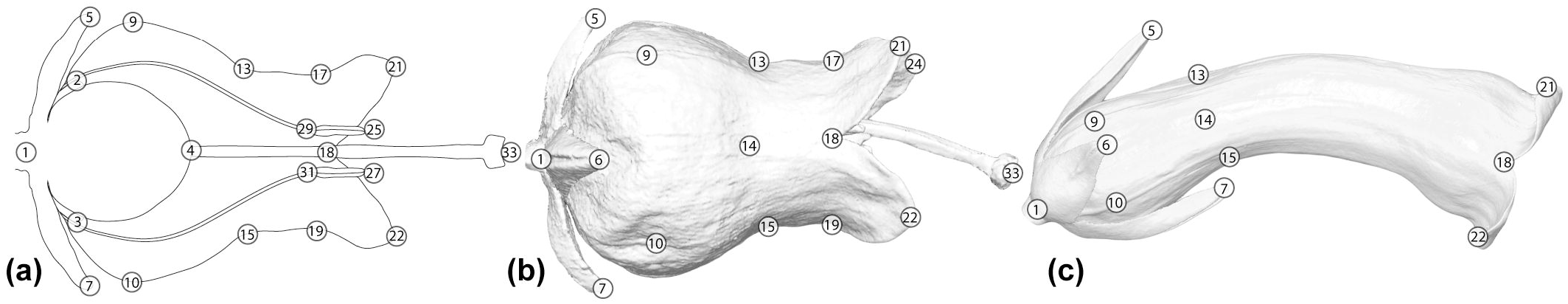
Landmarks. Landmarks used to digitise the shape of *Erica* flowers. (a) on schematic longitudinal section diagramme of an *Erica* flower. (b) on a 3D model of an actinomorphic flower (*E. hirtiflora*). (c) on a 3D model of a zygomorphic flower (*E. leucotrachela*).

### Pollination syndrome prediction

Given the scarcity of direct evidence for pollinators of particular *Erica* species, we relied on visitor data, combining published observations of populations in the wild (eight species, see table 1) with our own of individuals in cultivation (one species, see table 1), to assess pollination syndrome. Species with flowers observed to be visited by birds and long-proboscid flies (LPF) (see Table 1) were classified into the specialised bird and LPF syndrome. Wind pollination was documented in one species, which was then classified into the wind syndrome. Species with flowers that were observed to be visited by several groups of insects that could pollinate the flowers were classified into the generalist syndrome. Using these observations, we identified the floral shape and size components discriminating among pollination types using a random forests (RF) classification algorithm (Breiman, 2001). See Methods S1 for details.

### Modularity analysis

We used the RV coefficient method of Klingenberg (Klingenberg, 2009), implemented in MorphoJ (Klingenberg, 2011) to test our modularity hypotheses. The methodology uses the RV coefficient, a multivariate generalisation of the squared Pearson coefficient (Escoufier, 1973), as a measure of independence of subsets of the landmark data; it identifies sets of landmarks that group together and are likely to function as evolutionary entities. We carried out modularity analyses on subsets of our data pooled by syndrome (variation pooled by species). We then calculated the correlation between the shape variation of the sets of landmarks (RV coefficient) of the partitions corresponding to the *attraction-reproduction*, the developmental, and two different efficiency hypotheses (Fig. 1a-d, table S2) and compared it with that of 100 million random partitions. The proportion of partitions with lower RV coefficient than the tested partition (i.e. partitions showing higher among-set independence) was used as a measure of support for that partition, the lower the proportion, the higher the support (Young, 2006; Gomez *et al.*, 2014).

### Estimation of size, and integration

Size was measured as species-level average in centroid size, as implemented in MorphoJ. Integration coefficients were calculated at the species level as shape PCA eigenvalue variance scaled by the total variance and number of variables (Klingenberg & Marugan-Lobon, 2013) as implemented in MorphoJ (see Table S3).

### Phylogenetic inference

Phylogenetic relationships were inferred using DNA sequences from two loci of the chloroplast genome (*trn*LF-*ndhJ* and *trnT-L* intergenic spacers) and one loci of the nuclear genome (internal transcribed spacer (ITS)) from 61 pre-existing sequences of 19 *Erica* species as ingroup and *Calluna vulgaris* and *Daboecia cantabrica* as outgroups (see Table S4 for source of the sequences and their GenBank numbers). Divergence time analyses were carried out within a Bayesian framework by employing an uncorrelated lognormal relaxed clock model in BEAST version 1.8.4 (Drummond *et al.*, 2012) by applying secondary calibration via using the two previously published nodal ages (Pirie *et al.*, 2016). See Methods S1 for details.

### Ancestral Character State Reconstruction

We used a pruned phylogeny (i.e. removing the outgroup) for the 19 *Erica* species included in this study to estimate the probability of the pollination strategy states for all nodes of the phylogeny. As a demonstration of the potential of this approach, we estimated ancestral states of pollination syndromes using Maximum Likelihood (ML) (Harmon *et al.*, 2010; Revell, 2012) and empirical Bayes (Revell, 2012) methods. See Methods S1 and Table S5 for details.

### Models of floral trait evolution (unidimensional and high-dimensional)

We applied a penalised likelihood approach to high-dimensional phenotypic dataset of flower shapes of 19 *Erica* species to estimate the fit of three different evolutionary models; Brownian Motion (BM), Ornstein–Uhlenbeck (OU), and Early Burst (EB) in order to better understand the process of floral-shape evolution in the clade (Clavel *et al.*, 2018). The analysis was carried out under the *fit_t_pl* function (RPANDA)(Morlon *et al.*, 2016), and the best fit of the abovementioned three models was assessed using the Generalised Information Criterion (GIC) with the *GIC* function (mvmorph)(Clavel *et al.*, 2015). Finally, we employed the parameters derived from the evolutionary model that best fitted our high-dimensional data to obtain floral shape reconstructions through time, as implemented in the function *ancestral* and *phyl.pca_pl* (RPANDA)(Morlon et al., 2016). To visualise 3D models of the reconstructed ancestral floral shapes at selected nodes, a 3D surface model of a flower of *Erica hirtiflora* (lying approximatively in the middle of the PC1 x PC2 space plot) was warped to each target ancestral shape. This was carried out by aligning the reconstructed ancestral shape at the selected nodes and the landmark data of the chosen model (*E. hirtiflora*) using a thin plate spline (TPS) interpolation (Wiley et al., 2005), using the function *tps3d* (Morpho)(Schlager, 2017) and the function *extractShape* (Clavel *et al.*, 2018).

We fitted a series of likelihood models (i.e. Brownian motion and Ornstein-Uhlenbeck models) to understand how changes in pollination syndromes influence the evolution of various continuous unidimensional floral traits of *Erica* (i.e. PC1, PC2, PC3, PC4, PC5, centroid size, and integration). The best fitting model was determined comparing AICc, ΔAICc, and AICc weights among the models. All analyses were implemented using the R package OUwie (Beaulieu *et al.*, 2012). See Methods S1 for details.

## Results

### Pollination syndromes prediction

The floral features used in the Random Forest (RF) classification algorithm successfully classified species into pollinator classes. The most important variable for pollinator prediction was tube length (Fig. **S1a**, Tables S6, S7). The next 15 most important variables were landmarks describing the widest and narrowest positions of the corolla, the ovary/style transition, the meeting point of petal lobes, and the position of sepal tips (Fig. **S1a**, Table S6). For 9 of the 10 predicted species, all flowers were assigned to the same pollination syndrome (Table S8). *E. georgica* was classified either as generalist, bird, LPF, or wind syndrome with varying support (Table S8). We assigned *E*. *georgica* to the LPF syndrome because the tube length of all these flowers corresponds to that syndrome (Fig. **S1b**), and because the shape of the flower and its morphology also corresponds to that syndrome, as defined for *Erica* (Rebelo *et al.*, 1985). Our RF classifications are in agreement with (Rebelo *et al.*, 1985).

### Flower shape

Together, principal component (PC) 1 and PC2 account for 62 % of total shape variation (38.9% for PC1 and 22.1% for PC2). The main distortion along the PC1 is a constriction, elongation and slight curving of the corolla tube. Flowers along PC2 are mainly differentiated by the proximal to medial position of the inflation of the corolla. This varies from globose-urceolate to tubular-urceolate flowers along PC1 and cylindrical to ovoid floral shape along PC2. The PC axis-related distortion along PC1 and 2 is visualised by an exemplary shape distortion of a flower of *E. hirtiflora* (Fig. 3). The spreading along the two axes did not reflect the phylogeny in separating clades defined by (Pirie *et al.*, 2016) (but see the *Evolution* section below). The convergent evolution of bird and LPF syndrome in our dataset display different patterns: the unrelated LPF syndrome flowers are tightly clustered in the morphospace whereas the bird syndrome flowers are in two clusters.

**Figure 3.**
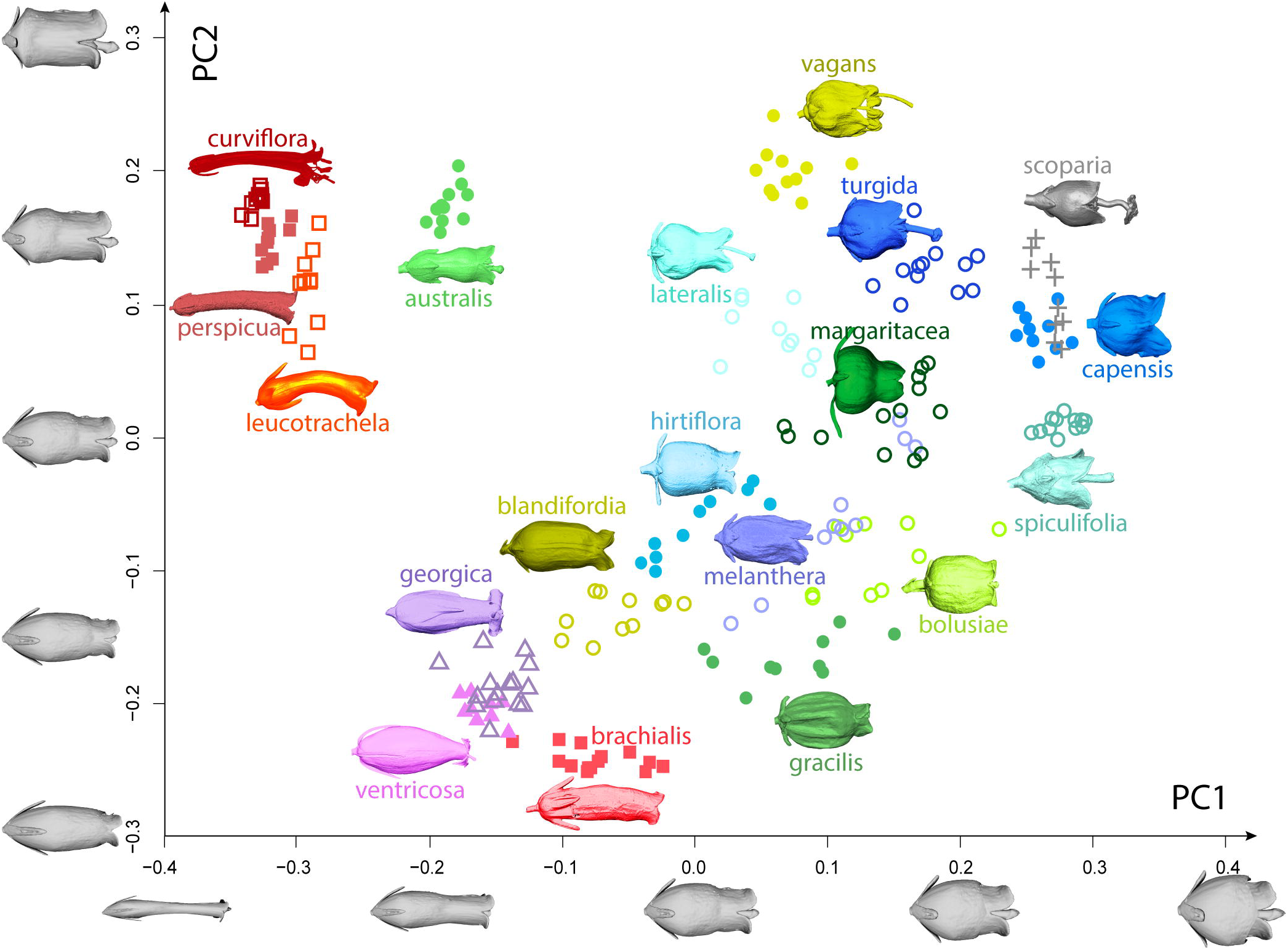
Shape PCA & syndromes. Two-dimensional ordination plot from a PCA analysis of 33 landmarks and 209 individual flowers of 19 *Erica* species. A representative flower surface-model for each species is plotted next to the dots corresponding to individual flowers of the same species. Colour and shape coding: green-blue circles, generalist syndrome, orange-red squares, bird syndrome, pink and purple triangles long-proboscid fly syndrome, grey crosses, wind syndrome. Closed symbols: observed visitors, open symbols: predicted visitors. Loadings of axes: x-axis PC1: 38.9% of shape variation, y-axis PC2: 22.1% of shape variation. In order to illustrate changes in floral shape associated with PC1 and PC2, a flower from the centre of the morphospace (*E. hirtiflora*) was distorted according to PC1 and PC2 and plotted along their respective axes.

### Modularity

In flowers with generalist syndrome, the best supported modularity hypothesis was the *developmental* hypothesis (see Table 2, Fig. 4a, Fig. **S2a-d**), although the *efficiency* hypotheses 1 and 2 received -weaker- support (see Table 2). In flowers with bird syndrome, the best supported modularity hypothesis was the *efficiency* 1 hypothesis (see Table 2, Fig. 4b, Fig. **S2e-h**), although the *developmental* hypothesis received –slightly weaker- support (see Table 2). In flowers with LPF syndrome, the best supported modularity hypothesis was the *efficiency* 2 hypothesis (see Table 2, Fig. 4c, Fig. **S2i-l**), although the *efficiency* hypothesis 1 received -weaker- support (see Table 2). In flowers with wind syndrome, the best supported modularity hypothesis was the *developmental* hypothesis (see Table 2, Fig. 4d, Fig. **S2m-p**). The *attraction-reproduction* hypothesis was not strongly supported for any pollination syndrome (See table 2).

**Table 2.** Modularity tests for the *attraction-reproduction*, *developmental*, and *efficiency 1* and *2* hypotheses. (Most significant values in bold).

| **Generalist syndrome** |  |  |  |
| --- | --- | --- | --- |
| hypothesis | RV of hypothesis | lowest RV | proportion lower RV |
| Attraction/reproduction | 0.22 | 0.19 | 1.40E-003 |
| **Developmental** | 0.12 | 0.11 | **2.30E-007** |
| Efficiency 1 | 0.13 | 0.11 | 4.10E-005 |
| Efficiency 2 | 0.16 | 0.14 | 7.33E-006 |

| **Bird syndrome** |  |  |  |
| --- | --- | --- | --- |
| hypothesis | RV of hypothesis | lowest RV | proportion lower RV |
| Attraction/reproduction | 0.4 | 0.16 | 3.50E-002 |
| Developmental | 0.19 | 0.16 | 3.66E-006 |
| **Efficiency 1** | 0.15 | 0.14 | **3.02E-006** |
| Efficiency 2 | 0.29 | 0.16 | 3.50E-003 |

| **LPF syndrome** |  |  |  |
| --- | --- | --- | --- |
| hypothesis | RV of hypothesis | lowest RV | proportion lower RV |
| Attraction/reproduction | 0.39 | 0.3 | 1.80E-002 |
| Developmental | 0.23 | 0.17 | 8.07E-004 |
| Efficiency 1 | 0.17 | 0.16 | 4.20E-006 |
| **Efficiency 2** | 0.23 | 0.22 | **5.50E-007** |

| **Wind syndrome** |  |  |  |
| --- | --- | --- | --- |
| hypothesis | RV of hypothesis | lowest RV | proportion lower RV |
| Attraction/reproduction | 0.72 | 0.44 | 2.50E-001 |
| **Developmental** | 0.43 | 0.32 | **1.70E-003** |
| Efficiency 1 | 0.47 | 0.29 | 4.00E-002 |
| Efficiency 2 | 0.54 | 0.34 | 2.60E-002 |

**Figure 4.**
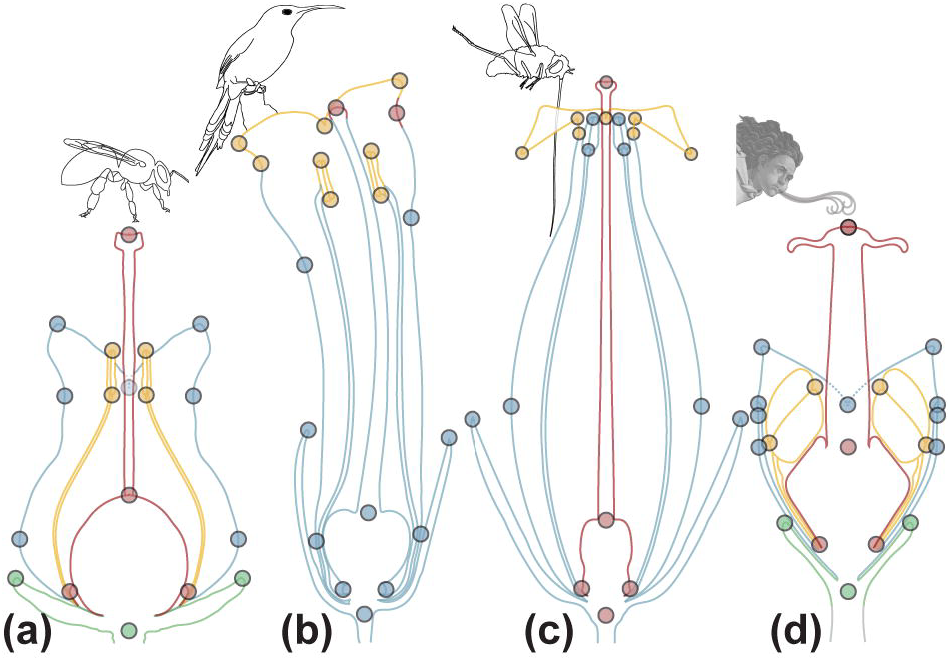
Modules in *Erica* flowers. (a) the best supported partition in flowers with generalist syndrome is the *developmental* hypothesis: a 4-fold partition with each organ class forms one module. (b) the best supported partition in the flowers with bird syndrome is the *efficiency* hypothesis 1, where the corolla lobes and the stamen form a putative “pollen deposition module”, and joining of the upper corolla lobes and the stigma form a putative “pollen receipt module”. The third set of landmarks comprises the rest of the flower. (c) the best supported partition in flowers with long-proboscid fly syndrome is the *efficiency* hypothesis 2, where the landmarks on the narrow corolla aperture form a putative “restriction module” that restricts access to the floral reward to only insects with very narrow proboscises. A second set of landmarks is formed by the gynoecium, and a third set of landmarks comprises the rest of the flower. (d) the best supported partition in flowers with wind syndrome the *developmental* hypothesis: a 4-fold partition with each organ class forms one module. Pollinator drawings, originals. Generalists represented by drawing of bee. Character representing the wind: Zephyr from “The birth of Venus” by Sandro Boticelli (ca. 1480).

### Allometry

The symmetric component of the entire dataset exhibited significant but weak allometry: 1.17% & (*P* = 0.001; see Fig. **S3a**). If the species are split by pollination syndrome, the proportion of variation explained by allometry (pooled by species) differs according to syndrome (see Notes S3). For the sake of brevity, only the allometric deformation in syndromes for which it is both strong (> 10% predicted shape) and significant (P < 0.05) will be discussed here (i.e. long-proboscid flies and wind syndromes). In the flowers with LPF syndrome, large flowers tend to have a more flask-shaped corolla, and the landmarks on the mouth of the corolla are closer to the floral axis (Fig. **S3b**). In the flowers with wind syndrome, large flowers tend to have corolla lobes more open and stamens more exerted (Fig. **S3c**).

### Ancestral Character States Reconstruction

Ancestral state reconstruction for pollination syndromes (Fig. 5a) suggests that the generalist pollination syndrome is the possible most recent common ancestral (MRCA) state in *Erica*. Within our sampled species the bird pollination syndrome, as well as the LPF syndrome evolved twice independently.

**Figure 5.**
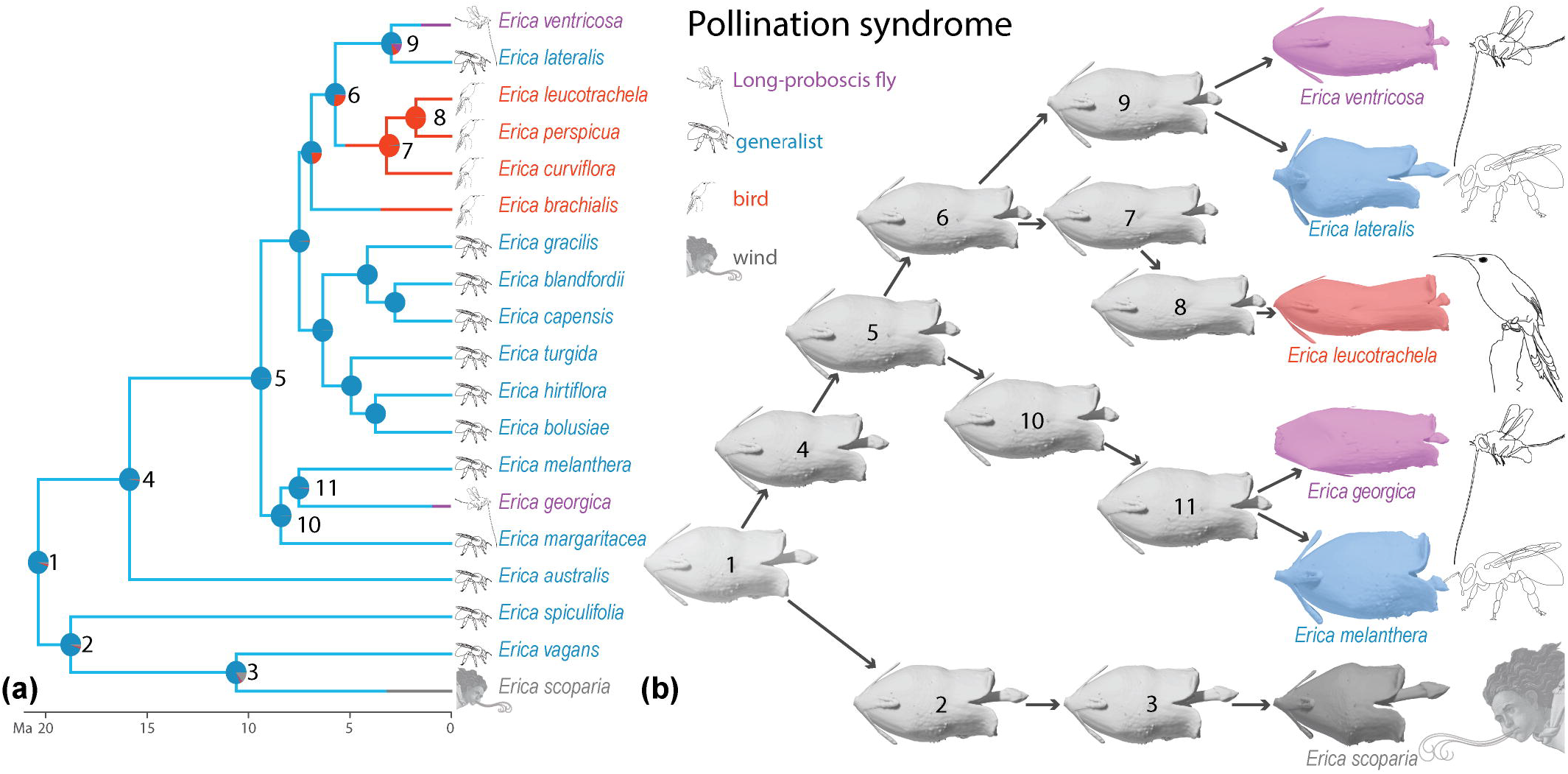
Ancestral state reconstruction for pollination syndromes and floral shape in *Erica*. (a) stochastic character mapping of the four pollination syndromes optimised on a chronogram inferred from Bayesian dating. Pie charts at internal nodes indicate the proportion of stochastic mapping from 1000 runs using the Equal Rates (ER) model. (b) ancestral shape reconstruction and reconstructed evolutionary trajectories for six selected species of *Erica*, including species from all four studied pollination syndromes and two convergent evolution of flowers with long-proboscid fly syndrome.

### Models of floral trait evolution

Under the penalised likelihood approach, the best fitting model to the evolution of the highly-dimensional whole floral shape in *Erica* was the Ornstein–Uhlenbeck model (OU; lowest GIC; Table S9), which assumes evolution towards an optimal floral shape mean as would be expected under selection.

The MRCA floral shape of *Erica* most likely displays short and urceolate flowers, as expected for flowers with generalist syndrome (Fig. 5b, node 1). The reconstructed evolutionary trajectory (under the best fitted model of OU) displays likely late differentiations in flower shape, with most differentiation possibly occurring at the most recent internal nodes of the tree (Fig. 5b, nodes 3, 7, 8, and 9). In both reconstructed ancestors of convergent evolution of LPF syndrome, the most recent internal nodes (Fig. 5b, nodes 9 and 11) likely display differentiation but this differentiation is weak compared to that of terminal nodes (Fig. 5b, flowers of *E. ventricosa*, and *E. georgica*).

The results of the fitting of five models (BM_1_, BM_S_, OU_1_, OU_M_, and OU_MV_) on quantitative floral trait evolution (shape PC1-5, size, and integration) under the four pollination-syndrome regimes are summarised in Table 3. The Hessian matrix of one model (i.e. OU_MV_) displayed a negative eigenvalue for PC3, PC4, integration, and centroid size, which means that this model was too complex for the information contained in these data and it was excluded from the analyses. Different evolutionary scenarios yielded variable AICc distributions, ΔAICc, and AICc weights (see Table 3). The evolution of floral shape along PC1 and centroid size of flowers were found to best fit an OU_M_ model (see Table 3). This evolutionary model suggests selection around four different optimal values (θ), one per pollination syndrome (see Table S10). This suggests that PC1 and centroid size have different evolutionary means for each of the four pollination syndrome regimes and that there is an evolutionary force that maintains PC1 and size closer to this evolutionary mean than would be expected under a BM model.

**Table 3.** Models of quantitative phenotypic trait evolution (PC1-5 of floral shape, size, and integration) under the pollination syndrome regime, and their biological interpretation, model fit of plausible models for the seven floral variables, indicating AICc (corrected AIC score), ΔAICc, and AICc weight..

| variables | Model | AICc | ΔAICc | AICc weight | Interpretation of the best model for shape, integration, and size variable evolution |
| --- | --- | --- | --- | --- | --- |
| PC1 | BM1 | -6.65 | 7.91 | 0.015 | Evolution of shape along PC1 is constrained; different optima depend on pollination syndromes, which would imply that optimal shape along PC1 has evolved separately for different pollination syndromes |
|  | BMS | 0.46 | 15.02 | 4.22E-04 |  |
|  | OU1 | -5.5 | 9.01 | 0.008 |  |
|  | **OUM** | **-14.56** | **0** | **0.772** |  |
|  | OUMV | -11.9 | 2.66 | 0.204 |  |
| PC2 | BM | -11.31 | 2.11 | 0.232 | Evolution of shape along PC2 is directed toward an optimum without being affected by the pollination syndromes |
|  | BMS | -4.76 | 8.66 | 0.009 |  |
|  | **OU1** | **-13.42** | **0** | **0.667** |  |
|  | OUM | -9.44 | 3.98 | 0.091 |  |
|  | OUMV | -1.27 | 12.16 | 0.002 |  |
| PC3 | **BM** | **-34.98** | **0** | **0.758** | Evolution of shape along PC3 is random and not affected by the different pollination syndromes |
|  | BMS | -26.25 | 8.73 | 0.010 |  |
|  | OU1 | -32.61 | 2.37 | 0.231 |  |
|  | OUM | -22.69 | 12.29 | 0.002 |  |
| PC4 | **BM** | **-38.35** | **0** | **0.630** | Evolution of shape along PC4 is random and not affected by the different pollination syndromes |
|  | BMS | -30.17 | 8.17 | 0.011 |  |
|  | OU1 | -37.03 | 1.32 | 0.326 |  |
|  | OUM | -32.5 | 5.85 | 0.034 |  |
| PC5 | BM | -37.62 | 0.4 | 0.437 | Evolution of shape along PC5 is directed toward an optimum without being affected by the pollination syndromes |
|  | BMS | -30.49 | 7.52 | 0.012 |  |
|  | **OU1** | **-38.01** | **0** | **0.533** |  |
|  | OUM | -31.22 | 6.8 | 0.018 |  |
|  | OUMV | -18.84 | 19.17 | 3.66E-05 |  |
| Integration | BM | -47.52 | 1.09 | 0.364 | Evolution of shape integration is directed toward an optimum without being affected by the pollination syndromes |
|  | BMS | -38.67 | 9.94 | 0.004 |  |
|  | **OU1** | **-48.61** | **0** | **0.6264** |  |
|  | OUM | -39.22 | 9.39 | 0.006 |  |
| Centroid size | BM1 | 160.28 | 23.11 | 9.42E-06 | Evolution of size is constrained; different optima depend on pollination syndromes, which would imply that optimal size has evolved separately for different pollination syndromes |
|  | BMS | 144.98 | 7.81 | 0.020 |  |
|  | OU1 | 161.09 | 23.91 | 6.29E-06 |  |
|  | **OUM** | **137.17** | **0** | **0.98** |  |

The evolution of floral shape along PC2, PC5, and floral integration were found to best fit an OU_1_ model (see Table 3). This result suggests that there is no difference between the four pollination syndromes, and that PC2, PC5, and integration each evolve towards a single one optimum (θ) across all *Erica* species (Table S10), indicating a lack of evidence for different constraints by the four pollination regimes. The best-fitted model for the evolution of floral shape along PC3 and PC4 was a BM_1_ model (see Table 3), where there is no difference between the pollination syndromes, and these floral variables evolve according to a random walk process.

## Discussion

### Modularity

In flowers with the generalist syndrome, our observation of strong support for the *developmental modularity* hypothesis supports the “trade-off” hypothesis of evolution of generalist flowers (which implies the absence of efficiency modules), and invalidates both the “trait specialisation” hypothesis and the “common shape” hypothesis (which both imply the evolution of efficiency modules). This contrasts with flowers with specialised syndromes (bird and LPF syndromes) which display support for (different) *efficiency* hypotheses. Similar patterns of modularity to that supported in flowers with bird syndrome (attraction-receipt-deposition) have been found across angiosperms: in a reanalysis of existing data, the least variable attributes of flowers were found to be those potentially affecting the mechanical fit between flower and pollinator (Cresswell, 1998). The results of (Cresswell, 1998) suggest independence, but do not test for the latter; such a test was carried out only in few studies such as for species of *Nicotiana* (Solanaceae) where such similar efficiency modules (lengths of the floral tube, stamens and gynoecium) were evidenced (Herrera *et al.*, 2002; Bissell & Diggle, 2010).

In the flowers with LFP syndrome, the set of landmarks of the “corolla aperture” does not include any reproductive organs; the function of this set is thus most likely not directly pollen deposition or receipt. In the Cape, flowers with LPF syndrome typically have very narrow floral tubes (Goldblatt & Manning, 2000); *Erica* flowers with this syndrome, however, do not always have narrow tubes, but do have a narrow corolla apertures (see Fig. 4c) (Rebelo *et al.*, 1985). This corolla aperture likely plays a role in restricting access to the floral rewards to certain classes of pollinators. This interpretation is supported by the allometric shape deformation (how shape changes with size) in LPF syndrome flowers: in shape, in larger flowers the corolla aperture is, relative to the rest of the flower, narrower, but in size, the corolla aperture stays about the same size in smaller and in larger flowers (Fig. **S3b**). Because of its putative function, we propose to refer to the set of landmarks on the corolla aperture as a “restriction module”. Similar structures were found to preclude visits from bats in bird-pollinated *Burmeistera* (Campanulaceae), and to vary much less than the rest of the flower (Muchhala, 2006), suggesting they constitute an independent module. Moreover, this restriction module also contains the petal tips (Fig. 4c), that do not actively contribute to limiting access to the floral reward; their small size relative to the rest of the corolla also precludes a major role in pollinator attraction. Their presence in the restriction module is therefore most likely non-adaptive and only due to their developmental proximity to the corolla aperture. Their presence within the restriction module is therefore most likely a spandrel *sensu* Gould and Lewontin (Gould & Lewontin, 1979). If it were feasible, a denser sampling of landmarks across the flowers would probably uncover more of such structures grouping in shape modules owing to their developmental proximity and not their function. In flowers with wind syndrome, support for the developmental hypothesis suggests that the shape of the different organ classes is independent form each other. This could be due to the fact that (1) wind pollinated flowers probably do not require across-organ class modules (for pollen receipt), and (2) that our data is dominated by developmental shape changes. Modelling studies in grasses that have shown that pollen deposition overwhelmingly relies only on direct impact on the stigma and not on air flows generated by the rest of the flower (Cresswell *et al.*, 2010), which suggests that there is no selection pressure for the rest of the flowers to form pollen receipt modules (as in *efficiency* hypothesis 1). The strong but weakly significant allometry reflects typical differences in flower shape related to differences in anthesis stage: larger (older) flowers have more open petals and more exerted stamens than smaller (younger) flowers (Fig. **S3c**); these changes would also cause organs classes to each display shape variation along their own developmental axis and be independent from each other. This notwithstanding, any interpretation is tentative given our limited sampling of this syndrome.

### Floral shape evolution

The radiation of *Erica* in the Cape is the greatest known to have occurred there and one of the greatest in recent plant biological history (Pirie *et al.*, 2016). Analyses confirmed the “hotbed” hypothesis in the genus, i.e. that the radiation of *Erica* was due to increased speciation rates, and showed an overall recent slowing down of speciation rates (although they do remain high in the former South Western clade (Pirie *et al.*, 2016)). Shifts in multiple local-scale ecological gradients, and repeated shift in pollinator preferences appear to have taken place (Linder *et al.*, 2010; Pirie *et al.*, 2016). Such a radiation fits Simpson’s adaptive zone model in which similar niches become ecologically available to a lineage, free from competitors (Simpson, 1944): when a lineage first enters these zones, phenotypical evolution should at first be fast, but as ecological niches are filled, the rate of phenotypical evolution should then slow down (Simpson, 1944; Schluter, 2000; Losos & Miles, 2002; Harmon *et al.*, 2010). In such a radiation, one would expect to recover an EB mode of phenotypical evolution (Harmon *et al.*, 2010). However, our analysis of the highly-dimensional morphometric dataset of flower shape recovered as the best fit an OU model of evolution (Table S9), a model considered to better represent the importance of selection. This is further supported by our ancestral floral shape reconstruction (Fig. 5b), which indicates a pattern of greater phenotypical variation at the most recent internal nodes of the tree (Figs. 5a & b, nodes 3, 8, 9, 11), a pattern consistent with pollinator-driven selection (OU model (Harmon *et al.*, 2010)). Our finding of strong evolutionary changes over short time scales concurs with previous findings from diverse data sources (Gingerich, 1983; Lynch, 1990; Hendry & Kinnison, 1999; Roopnarine, 2003; Estes & Arnold, 2007; Harmon *et al.*, 2010). This is furthermore strongly supported by our analyses of the evolutionary model of PC1 and centroid size under different regimes (i.e. pollination syndrome) which recovered as best fit an OU_M_ model of evolution (selection towards different optima; Tables 3, S11), strongly indicating that pollinators have indeed driven the evolution of floral shape (see below), therefore supporting the a strong role for pollinator-driven speciation in *Erica* (Pirie *et al.*, 2011). PC1 corresponds to a shape change from open bell shaped flowers to more elongated, tubular flowers, generating, for the same size, longer tubes and strongly affecting the landmarks on the narrowest and broadest parts of the corolla (see Fig. 3). These landmarks, together with tube length, were shown by our random forest analyses to be especially important in predicting pollination syndromes (see Table S6). PC1 therefore involves a shape change that is especially relevant for the generation of the different floral shapes of the different pollination syndromes. Variation in PC1 was thus most likely co-opted by evolution to generate the different syndrome morphologies, and ended up encapsulating almost 40% of shape variance (Table S10). Similarly, centroid size is strongly correlated with tube length (R^2^ = 0.96; P= 2.2E-16), the variable we demonstrate to play the strongest role in predicting the different syndromes (Fig. **S1a**, Table S6). Other PCs probably do not generate variation for which divergent selection on syndromes was present (or strong enough to be identified with our limited sampling), and therefore follow either a single optimum (OU_1_) or a random model (BM_1_) of evolution (Tables 3, S11).

Our result, that integration follows an OU_1_ model of evolution (selection with a single optimum; Tables 3, S11), does not support increased floral integration in specialist compared to generalist flowers. Our results also contrast with the results of Gomez et al. (2014) who recovered a BM model of evolution for floral integration (Gomez *et al.*, 2014). However, Gomez et al. (2014) included only landmarks placed on the petals (in 2D), whereas our study includes reproductive organs (in 3D). Because, efficiency modularity (including reproductive parts) has been shown to be stronger than attraction modularity (including the petals only) (Rosas-Guerrero *et al.*, 2011), our study likely includes a signal that is not present in that of Gomez et al. (2014). Evolution of whole-flower integration towards a single optimum suggest that evolution of increased integration in functional part of the flowers may come at the cost of lower integration with other parts of the flowers, leading to evolution towards a single optimal value in generalised and specialised systems. Our findings thus do not support changes in integration as a whole, but strongly support changes in its structure, an observation congruent with (Ordano *et al.*, 2008).

## Conclusion

Our results illustrate for the first time the potential of 3D datasets (that include the reproductive organs of flowers) together with geometric morphometrics to uncover the modularity of the highly dimensional shape of flowers as a function of pollinator syndrome, and together with a novel penalised likelihood framework (Clavel *et al.*, 2018) also for the first time to test the fits of evolutionary models to the macro-evolution of high-dimensional flower shape and reconstruct its trajectory.

Simulations of biological evolution have demonstrated that modularity is favoured within environments where selection changes over time in such a way that each new selective pressure shares some of the aspects of the previous selective pressure (Kashtan & Alon, 2005; Kashtan *et al.*, 2007). It has been shown that within a pollination syndrome, selection on floral traits can change from year to year due to fluctuations in pollinator abundance (Herrera, CM, 1988; Campbell, 1989; Campbell *et al.*, 1991). We thus speculate that syndromes are such a changing environment, that evolution of a new syndrome is the equivalent to a change of environment, necessitating the evolution of a new modular organisation (although overall floral integration need not change), and finally that fluctuations in pollinator abundance (within a syndrome) play a role in the emergence of flower modularity.

## Acknowledgements

The authors wish to thank the Belvedere garden for granting access to the material (special thanks to the head gardener, Mr. M. Knaack). We would like to also thank the company Stadler Fleurs for donating material for this study. We also thank Julien Clavel for his help with the analyses, and Michael Pirie for his valuable comments on a previous version of the manuscript.

## Author contributions

YS and MvB designed the project, YS and MvB collected the material, DR and AB collected the data and ran preliminary analyses, MC carried out the random forest analyses, SM and MS carried out the phylogenetic analyses, SM carried out the trait evolution analyses, YS and CK carried out the geometric morphometric analyses, SM designed the manuscript, YS and SM wrote the manuscript. DR and AB contributed equally.

---

**Figure S1 Machine learning.** (a) landmark coordinates and tube length sorted by mean accuracy decrease in predicting pollination syndrome via Random Forest (the tube length is the best variable to predict pollination syndrome). (b) tube length (in mm) in studied species.

**Figure S2 Hypotheses test: RV distributions.** X-axis RV coefficient, y-axis frequency of values. Red arrow indicates the value of the RV coefficient of the modularity hypothesis tested. Left, hypotheses tested, right results of test. a-d tests for species with generalist syndrome. a, test of *attraction-reproduction* hypothesis. b, test for *efficiency* hypothesis 1. c, test for *efficiency* hypothesis 2. d, test for *developmental* hypothesis. e-h tests for species with bird syndrome. e, test of *attraction-reproduction* hypothesis. f, test for *efficiency* hypothesis 1. g, test for *efficiency* hypothesis 2. h, test for *developmental* hypothesis. i-l tests for species with long-proboscid fly syndrome. i, test of *attraction-reproduction* hypothesis. j, test for *efficiency* hypothesis 1. k, test for *efficiency* hypothesis 2. l, test for *developmental* hypothesis. m-p tests for species with wind syndrome. m, test of *attraction-reproduction* hypothesis. n, test for *efficiency* hypothesis 1. o, test for *efficiency* hypothesis 2. p, test for *developmental* hypothesis. Pollinator drawings, originals. Generalists represented by drawing of bee. Character representing the wind: Zephyr from “The birth of Venus” by Sandro Boticelli (ca. 1480).

**Figure S3 Allometry in *Erica* flowers.** (a) allometric plot. x-axis, log centroid size, y-axis shape axis. All 209 individual flowers from all 19 species studied are plotted. Blue to green dots generalist syndrome, orange to red squares bird syndrome, pink and purple triangles long-proboscid fly syndrome, gray crosses wind syndrome. (b) allometric deformation in flowers with long-proboscid syndrome for a change in log centroid size of 0.2. Pink, schematic drawing of smaller flowers, blue schematic drawing of larger flowers. (c) allometric deformation in flowers with wind syndrome for a change in log centroid size of 0.2. Pink, schematic drawing of smaller flowers, blue schematic drawing of larger flowers.

## List of supplementary data

**Table S1.** Species, sample numbers (n) and scanning conditions of *Erica* flowers.

**Table S2.** Landmarks used to digitise the shape of *Erica* flowers, and modules to which they belong in the modularity hypotheses tested.

**Table S3.** Species-level average values for size and integration.

**Table S4.** Genbank accession numbers for nrDNA ITS and cpDNA trnL-F-ndhJ and trnT-L sequence data.

**Table S5.** Discrete character mapping models for pollination syndromes.

**Table S6.** Main variables mean accuracy decrease of random forest syndrome prediction.

**Table S7.** Corolla tube length per flower.

**Table S8.** Classification of 114 individual flowers of diverse *Erica* species into the pollination syndromes,

**Table S9.** Support values evolutionary models of floral shape evolution.

**Table S10.** Summary of the preferred models of evolution for seven phenotypic trait variables (PC1-5 of floral shape, centroid size, and integration

**Methods S1.** This file contains details of the methodology used to for: X-ray tomography, 3D-landmarking, geometric morphometrics, pollination syndrome prediction, modularity analyses (exploratory and confirmatory approaches), phylogenetic inference, ancestral character states reconstruction, and models of trait evolution.

**Notes S1.** Literature analysis.

**Notes S2.** Allometric regressions and correlation between the corolla tube length and centroid size

**Scan data.** All the scan data will be deposited on PHAIDRA, the open data repository of the University of Vienna (https://phaidra.univie.ac.at/).

## References

Aigner PA. 2001. Optimality modeling and fitness trade□offs: when should plants become pollinator specialists? Oikos 95(1): 177–184.

Aigner PA. 2006. The evolution of specialized floral phenotypes in a fine-grained pollination environment. Plant–pollinator interactions: From specialization to generalization 23: 46.

Alon U. 2003. Biological networks: the tinkerer as an engineer. Science 301(5641): 1866–1867.

Arendse B. 2015. Variation in breeding systems and consequences for reproductive traits in Erica. University of Cape Town.

Armbruster WS, Tuxill JD, Flores TC, Vela JL. 1999. Covariance and decoupling of floral and vegetative traits in nine Neotropical plants: a re□evaluation of Berg’s correlation[pleiades concept. American Journal of Botany 86(1): 39–55.

Armbruster WS, Wege JA. 2018. Detecting canalization and intra-floral modularity in triggerplant (Stylidium) flowers: correlations are only part of the story. Annals of Botany 123(2): 355–372.

Bardua C, Wilkinson M, Gower DJ, Sherratt E, Goswami A. 2019. Morphological evolution and modularity of the caecilian skull. BMC evolutionary biology 19(1): 30.

Beaulieu JM, Jhwueng DC, Boettiger C, O’Meara BC. 2012. Modeling stabilizing selection: expanding the Ornstein–Uhlenbeck model of adaptive evolution. Evolution: International Journal of Organic Evolution 66(8): 2369–2383.

Bell G. 1997. The basics of selection: Springer Science & Business Media.

Berg RL. 1960. The ecological significance of correlation pleiades. Evolution 14(2): 171–180.

Bissell E, Diggle P. 2010. Modular genetic architecture of floral morphology in Nicotiana: quantitative genetic and comparative phenotypic approaches to floral integration. Journal of evolutionary biology 23(8): 1744–1758.

Bissell EK, Diggle PK. 2008. Floral morphology in Nicotiana: architectural and temporal effects on phenotypic integration. International Journal of Plant Sciences 169(2): 225–240.

Breiman L. 2001. Random forests. Machine learning 45(1): 5–32.

Campbell DR. 1989. Measurements of selection in a hermaphroditic plant: variation in male and female pollination success. Evolution 43(2): 318–334.

Campbell DR, Waser NM, Price MV, Lynch EA, Mitchell RJ. 1991. Components of phenotypic selection: pollen export and flower corolla width in Ipomopsis aggregata. Evolution 45(6): 1458–1467.

Carvallo G, Medel R. 2005. The modular structure of the floral phenotype in Mimulus luteus var. luteus (Phrymaceae). Revista Chilena de Historia Natural 78(4).

Chazot N, Panara S, Zilbermann N, Blandin P, Le Poul Y, Cornette R, Elias M, Debat V. 2016. Morpho morphometrics: shared ancestry and selection drive the evolution of wing size and shape in Morpho butterflies. Evolution 70(1): 181–194.

Clavel J, Aristide L, Morlon H. 2018. A Penalized Likelihood framework for high-dimensional phylogenetic comparative methods and an application to new-world monkeys brain evolution. Systematic Biology 68(1): 93–116.

Clavel J, Escarguel G, Merceron G. 2015. mvMORPH: an R package for fitting multivariate evolutionary models to morphometric data. Methods in Ecology and Evolution 6(11): 1311–1319.

Cresswell J. 1998. Stabilizing selection and the structural variability of flowers within species. Annals of Botany 81(4): 463–473.

Cresswell JE, Krick J, Patrick MA, Lahoubi M. 2010. The aerodynamics and efficiency of wind pollination in grasses. Functional Ecology 24(4): 706–713.

Cullinan J, Strenberg K, Tribe G. Simon’s Town, Western Cape, South Africa: Ujubee.

Diggle PK. 2014. Modularity and intra-floral integration in metameric organisms: plants are more than the sum of their parts. Phil. Trans. R. Soc. B 369(1649): 20130253.

Drake AG, Klingenberg CP. 2010. Large-scale diversification of skull shape in domestic dogs: disparity and modularity. The American Naturalist 175(3): 289–301.

Drummond AJ, Suchard MA, Xie D, Rambaut A. 2012. Bayesian phylogenetics with BEAUti and the BEAST 1.7. Molecular biology and evolution 29(8): 1969–1973.

Edwards C, Weinig C. 2011. The quantitative-genetic and QTL architecture of trait integration and modularity in Brassica rapa across simulated seasonal settings. Heredity 106(4): 661.

Ellis AG, Brockington SF, de Jager ML, Mellers G, Walker RH, Glover BJ. 2014. Floral trait variation and integration as a function of sexual deception in Gorteria diffusa. Phil. Trans. R. Soc. B 369(1649): 20130563.

Escoufier Y. 1973. Le traitement des variables vectorielles. Biometrics: 751–760.

Estes S, Arnold SJ. 2007. Resolving the paradox of stasis: models with stabilizing selection explain evolutionary divergence on all timescales. The American Naturalist 169(2): 227–244.

Esteve□Altava B. 2017. In search of morphological modules: a systematic review. Biological Reviews 92(3): 1332–1347.

Fabbri M, Koch NM, Pritchard AC, Hanson M, Hoffman E, Bever GS, Balanoff AM, Morris ZS, Field DJ, Camacho J. 2017. The skull roof tracks the brain during the evolution and development of reptiles including birds. Nature ecology & evolution 1(10): 1543.

Faegri K, Van Der Pijl L. 1966. Principles of pollination ecology. Oxford, UK: Pergman Press.

Fenster CB, Armbruster WS, Wilson P, Dudash MR, Thomson JD. 2004. Pollination syndromes and floral specialization. Annu. Rev. Ecol. Evol. Syst. 35: 375–403.

Fern K, Fern A 2012. Erica vagans - L. U.K.: Plants For A Future (Company No. 3204567, Charity No. 1057719).

Fornoni J, Ordano M, Pérez-Ishiwara R, Boege K, Domínguez CA. 2015. A comparison of floral integration between selfing and outcrossing species: a meta-analysis. Annals of Botany 117(2): 299–306.

Geerts S. 2011. Assembly and disassembly of bird pollination communities at the Cape of Africa. Stellenbosch: Stellenbosch University.

Gil-López MJ, Segarra-Moragues JG, Ojeda F. 2014. Population genetic structure of a sandstone specialist and a generalist heath species at two levels of sandstone patchiness across the Strait of Gibraltar. PloS one 9(5): e98602.

Gingerich P. 1983. Rates of evolution: effects of time and temporal scaling. Science 222: 159–162.

Goldblatt P, Manning JC. 2000. The long-proboscid fly pollination system in southern Africa. Annals of the Missouri Botanical Garden: 146–170.

Gomez JM, Perfectti F, Klingenberg CP. 2014. The role of pollinator diversity in the evolution of corolla-shape integration in a pollination-generalist plant clade. Philosophical Transactions of the Royal Society B-Biological Sciences 369(1649).

Gomez JM, Torices R, Lorite J, Klingenberg CP, Perfectti F. 2016. The role of pollinators in the evolution of corolla shape variation, disparity and integration in a highly diversified plant family with a conserved floral bauplan. Annals of Botany 117(5): 889–904.

González AV, Murúa MM, Pérez F. 2015. Floral integration and pollinator diversity in the generalized plant-pollinator system of Alstroemeria ligtu (Alstroemeriaceae). Evolutionary ecology 29(1): 63–75.

Gould SJ, Lewontin RC. 1979. The spandrels of San Marco and the Panglossian paradigm: a critique of the adaptationist programme. Proceedings of the Royal Society of London. Series B. Biological Sciences 205(1161): 581–598.

Grant V, Grant KA. 1965. Flower pollination in the Phlox family.

Hansen TF, Carter AJ, Pélabon C. 2006. On adaptive accuracy and precision in natural populations. The American Naturalist 168(2): 168–181.

Harmon LJ, Losos JB, Jonathan Davies T, Gillespie RG, Gittleman JL, Bryan Jennings W, Kozak KH, McPeek MA, Moreno□Roark F, Near TJ. 2010. Early bursts of body size and shape evolution are rare in comparative data. Evolution: International Journal of Organic Evolution 64(8): 2385–2396.

Hendry AP, Kinnison MT. 1999. Perspective: the pace of modern life: measuring rates of contemporary microevolution. Evolution 53(6): 1637–1653.

Herrera CM. 1988. Variation in mutualisms: the spatiotemporal mosaic of a pollinator assemblage. Biological Journal of the Linnean Society 35(2): 95–125.

Herrera CM. 2001. Deconstructing a floral phenotype: do pollinators select for corolla integration in Lavandula latifolia? Journal of evolutionary biology 14(4): 574–584.

Herrera CM 2009. Multiplicity in unity: plant subindividual variation and interactions with animals. University of Chicago Press, 1–4.

Herrera CM, Cerdá X, Garcia M, Guitián J, Medrano M, Rey PJ, Sánchez□Lafuente A. 2002. Floral integration, phenotypic covariance structure and pollinator variation in bumblebee□pollinated Helleborus foetidus. Journal of evolutionary biology 15(1): 108–121.

Herrera J. 1988. Pollination relationships in southern Spanish Mediterranean shrublands. The Journal of Ecology: 274–287.

Heystek A, Geerts S, Barnard P, Pauw A. 2014. Pink flower preference in sunbirds does not translate into plant fitness differences in a polymorphic Erica species. Evolutionary ecology 28(3): 457–470.

Heywood JS, Michalski JS, McCann BK, Russo AD, Andres KJ, Hall AR, Middleton TC. 2017. Genetic and environmental integration of the hawkmoth pollination syndrome in Ruellia humilis (Acanthaceae). Annals of Botany 119(7): 1143–1155.

Hulsey CD, de León FG, Rodiles□Hernández R. 2006. Micro□and macroevolutionary decoupling of cichlid jaws: a test of Liem’s key innovation hypothesis. Evolution 60(10): 2096–2109.

Johnson SD. 2006. Pollinator-driven speciation in plants. Ecology and evolution of flowers: 295–310.

Johnson SD, Steiner KE. 2000. Generalization versus specialization in plant pollination systems. Trends in ecology & evolution 15(4): 140–143.

Joly S, Lambert F, Alexandre H, Clavel J, Léveillé□Bourret É, Clark JL. 2018. Greater pollination generalization is not associated with reduced constraints on corolla shape in Antillean plants. Evolution 72(2): 244–260.

Kashtan N, Alon U. 2005. Spontaneous evolution of modularity and network motifs. Proceedings of the National Academy of Sciences 102(39): 13773–13778.

Kashtan N, Noor E, Alon U. 2007. Varying environments can speed up evolution. Proceedings of the National Academy of Sciences 104(34): 13711–13716.

Kelber A, Jacobs GH 2016. Evolution of color vision. Human Color Vision: Springer, 317–354.

Kirsten H, Hogeweg P. 2011. Evolution of networks for body plan patterning; interplay of modularity, robustness and evolvability. PLoS computational biology 7(10): e1002208.

Klingenberg CP. 2009. Morphometric integration and modularity in configurations of landmarks: tools for evaluating a priori hypotheses. Evolution & development 11(4): 405–421.

Klingenberg CP. 2011. MorphoJ: an integrated software package for geometric morphometrics. Molecular Ecology Resources 11: 353–357.

Klingenberg CP. 2014. Studying morphological integration and modularity at multiple levels: concepts and analysis. Philosophical Transactions of the Royal Society B-Biological Sciences 369(1649).

Klingenberg CP, Debat V, Roff DA. 2010. Quantitative genetics of shape in cricket wings: developmental integration in a functional structure. Evolution: International Journal of Organic Evolution 64(10): 2935–2951.

Klingenberg CP, Marugan-Lobon J. 2013. Evolutionary Covariation in Geometric Morphometric Data: Analyzing Integration, Modularity, and Allometry in a Phylogenetic Context. Systematic Biology 62(4): 591–610.

Linder HP, Johnson SD, Kuhlmann M, Matthee CA, Nyffeler R, Swartz ER. 2010. Biotic diversity in the Southern African winter-rainfall region. Current opinion in environmental sustainability 2(1-2): 109–116.

Losos JB, Miles DB. 2002. Testing the hypothesis that a clade has adaptively radiated: iguanid lizard clades as a case study. The American Naturalist 160(2): 147–157.

Lynch M. 1990. The rate of morphological evolution in mammals from the standpoint of the neutral expectation. The American Naturalist 136(6): 727–741.

McAdams HH, Srinivasan B, Arkin AP. 2004. The evolution of genetic regulatory systems in bacteria. Nature Reviews Genetics 5(3): 169–178.

Meng JL, Zhou XH, Zhao ZG, Du GZ. 2008. Covariance of floral and vegetative traits in four species of Ranunculaceae: a comparison between specialized and generalized pollination systems. Journal of integrative plant biology 50(9): 1161–1170.

Morlon H, Lewitus E, Condamine FL, Manceau M, Clavel J, Drury J. 2016. RPANDA: an R package for macroevolutionary analyses on phylogenetic trees. Methods in Ecology and Evolution 7(5): 589–597.

Morris JL, Puttick MN, Clark JW, Edwards D, Kenrick P, Pressel S, Wellman CH, Yang Z, Schneider H, Donoghue PCJ. 2018. The timescale of early land plant evolution. Proc Natl Acad Sci U S A 115(10): E2274–E2283.

Muchhala N. 2006. The pollination biology of Burmeistera (Campanulaceae): specialization and syndromes. American Journal of Botany 93(8): 1081–1089.

Notten A 2012. Erica brachialis Salisb. In PlantZAfrica, Institute SANB. South Africa: Kirstenbosch National Botanical Garden.

Ordano M, Fornoni J, Boege K, Domínguez CA. 2008. The adaptive value of phenotypic floral integration. New Phytologist 179(4): 1183–1192.

Pérez-Barrales R, Simón-Porcar VI, Santos-Gally R, Arroyo J. 2014. Phenotypic integration in style dimorphic daffodils (Narcissus, Amaryllidaceae) with different pollinators. Phil. Trans. R. Soc. B 369(1649): 20130258.

Pérez□Barrales R, Arroyo J, Scott Armbruster W. 2007. Differences in pollinator faunas may generate geographic differences in floral morphology and integration in Narcissus papyraceus (Amaryllidaceae). Oikos 116(11): 1904–1918.

Pérez F, Arroyo MT, Medel R. 2007. Phylogenetic analysis of floral integration in Schizanthus (Solanaceae): does pollination truly integrate corolla traits? Journal of evolutionary biology 20(5): 1730–1738.

Pigliucci M, Preston K. 2004. Phenotypic integration: studying the ecology and evolution of complex phenotypes: Oxford University Press.

Pirie M, Oliver E, De Kuppler AM, Gehrke B, Le Maitre N, Kandziora M, Bellstedt D. 2016. The biodiversity hotspot as evolutionary hot-bed: spectacular radiation of Erica in the Cape Floristic Region. BMC evolutionary biology 16(1): 190.

Pirie MD, Oliver E, Bellstedt DU. 2011. A densely sampled ITS phylogeny of the Cape flagship genus Erica L. suggests numerous shifts in floral macro-morphology. Molecular phylogenetics and evolution 61(2): 593–601.

Plants_Database 2019. Cornish Heath (Erica vagans). In Association NG. U.S.A.

Rebelo A, Siegfried W, Crowe A. 1984. Avian pollinators and the pollination syndromes of selected mountain fynbos plants. South African Journal of Botany 3(5): 285–296.

Rebelo A, Siegfried W, Oliver E. 1985. Pollination syndromes of Erica species in the south-western Cape. South African Journal of Botany 51(4): 270–280.

Revell LJ. 2012. phytools: an R package for phylogenetic comparative biology (and other things). Methods in Ecology and Evolution 3(2): 217–223.

Roopnarine PD. 2003. Analysis of rates of morphologic evolution. Annual Review of Ecology, Evolution, and Systematics 34(1): 605–632.

Rosas□Guerrero V, Quesada M, Armbruster WS, Pérez□Barrales R, Smith SD. 2011. Influence of pollination specialization and breeding system on floral integration and phenotypic variation in Ipomoea. Evolution: International Journal of Organic Evolution 65(2): 350–364.

Sahli HF, Conner JK. 2011. Testing for conflicting and nonadditive selection: floral adaptation to multiple pollinators through male and female fitness. Evolution: International Journal of Organic Evolution 65(5): 1457–1473.

Schlager S 2017. Morpho and Rvcg–Shape Analysis in R: R-Packages for geometric morphometrics, shape analysis and surface manipulations. Statistical shape and deformation analysis: Elsevier, 217–256.

Schluter D. 2000. The ecology of adaptive radiation: OUP Oxford.

Simpson GG. 1944. Tempo and mode in evolution: Columbia University Press.

Staedler YM, Masson D, Schönenberger J. 2013. Plant tissues in 3D via X-ray tomography: simple contrasting methods allow high resolution imaging. PloS one 8(9): e75295.

Stebbins GL. 1950. Variation and evolution in plants: Geoffrey Cumberlege.; London.

Stebbins GL. 1970. Adaptive radiation of reproductive characteristics in angiosperms, I: pollination mechanisms. Annual Review of Ecology and Systematics 1(1): 307–326.

Stevens P, Luteyn J, Oliver E, Bell T, Brown E, Crowden R, George A, Jordan G, Ladd P, Lemson K 2004. Ericaceae. Flowering Plants· Dicotyledons: Springer, 145–194.

Theophrastus ToE, Hort SA 1916. Of the parts of plants and their composition. Of classification. Inquiry Into Plants and Minor Works on Odours and Weather Signs: Heinemann.

Turner R 2010. Sunbird Surprise! Surprises in the world of plant pollination. Full Circle Magazine. U.K.: Full Circle Team. 34.

Vogel S 1954. Bliitenbiologische Typen als Elemente der Sippengliederung: Fischer—Jena.

Wagner GP, Altenberg L. 1996. Perspective: complex adaptations and the evolution of evolvability. Evolution 50(3): 967–976.

Wagner GP, Pavlicev M, Cheverud JM. 2007. The road to modularity. Nature Reviews Genetics 8(12): 921–931.

Weismann A. 1892. Das Keimplasma: eine theorie der Vererbung: Fischer.

Wiley DF, Amenta N, Alcantara DA, Ghosh D, Kil YJ, Delson E, Harcourt-Smith W, Rohlf FJ, St John K, Hamann B 2005. Evolutionary morphing. VIS 05. IEEE Visualization, 2005.: IEEE. 431–438.

Young NM. 2006. Function, ontogeny and canalization of shape variance in the primate scapula. Journal of Anatomy 209(5): 623–636.

