## Supplementary material for "Modularity and evolution of flower shape: the role of efficiency, development, and spandrels in *Erica*": Methods S1 (supplementary methods)

*Plant material*

The plant material was collected in 2016 and 2017, fixed in FAA, and stored at 4°C. Vouchers were deposited at the herbarium of the University of Vienna (WU). Flowers were collected from 1-3 individual(s) per species. Individuals of the same species were the product of clonal reproduction (which is the standard procedure in *Erica* cultivation) and common garden cultivation; moreover, the shape from flowers from clones of the same genotype did not display significant differentiation (data not shown). The material was thus treated as coming from a single individual per species.

*X-ray tomography*

For X-ray computed tomography, fixed flowers were transferred in 1% phosphotungstic acid in 70% EtOH and infiltrated for at least one week in order to increase contrast (Staedler *et al.*, 2013). Ten–fifteen individual flowers per species were batch-mounted in acryl pillow foam (DACRON® COMFOREL® INVISTA Ltd.) in cylindrical containers (VWR International, polypropylene) in EtOH vapour and sealed with PARAFILM (Staedler *et al.*, 2013). Scans were performed using a Zeiss Microscopy XCT-200 scanner. Scanning conditions are summarised in Table S1. The 209 flowers were scanned and used for landmarking. The raw scanning data was processed with the XMReconstructor package and 3D rendering of individual flowers was carried out in AMIRA 5.4.1 (Visualisation Sciences Group, SAS).

*3D-landmarking & Geometric Morphometrics*

For the sake of homogeneity between actinomorphic and zygomorphic flowers, we only used one symmetry plane to symmetrise our data. For calculation of the shape covariance matrix, we pooled variance within species and carried out downstream analyses only on the symmetrical component of shape variation. For syndrome-level analyses, we subdivided our dataset into groups of species with the same syndrome (variation pooled by species) and carried out the analyses on these sub-pools. Integration values, per species, were obtained by scaling the PCA eigenvalue variance by the total variance and the number of variables (Young, 2006; Klingenberg & Marugan-Lobon, 2013). PCA has been shown to be consistent in cases where few eigenvalues dominate a population covariance matrix (Jung & Marron, 2009), which is the case in our datasets (PC1–PC3 explain 72% of total variance in pooled dataset for all species); we therefore consider our values plausible estimates despite our restricted species-level sampling. Allometric regressions were carried out in MorphoJ on the symmetric component of shape variation (variation pooled by species).

*Pollination syndrome prediction*

In order to predict pollination syndromes, we used the function *randomForest* (randomForest)(Liaw & Wiener, 2002). We drew 500 forests of 1,001 trees each, with a number of variables sampled for splitting at each node (*mtry*) set to 4 (following (Liaw & Wiener, 2002) and (Breiman, 2003)). These trees were trained to discriminate among the different types of pollination based on the morphological variables "tube length" plus 99 procrustes coordinates describing the 3D shape of each flower. Differences among pollination syndromes were assessed for each discriminating variable with Kruskal-Wallis tests using *kruskal.test* (stat) (R_Core_Team, 2018) and *kruskalmc* (pgirmess) (Giraudoux, 2017) as post hoc tests, with a Bonferroni correction for multiple comparison tests. To assign a pollination syndrome to *Erica* species with unknown pollinator, we used the 500 trained forests from the previous RF classification and the function *predict* (*stat*) (R_Core_Team, 2018).

*Phylogenetic inference*

The molecular dating analysis was based on a secondary calibration from a previous study (Pirie *et al.*, 2016). Calibrations using the two previously published nodal ages were modelled with a normal distribution, where 95% of the prior weight fell within the highest posterior density (HPD) interval in which each node was discovered in the original study. Following (Pirie *et al.*, 2016), the crown node of the genus *Erica* was calibrated using a normal distribution prior with a median=22.5 Mya (StDev= 2.3), and the stem node of the genus (i.e. root of the tree) was calibrated using a normal distribution with a median=60.5 Mya (StDev= 7.9). After selecting a GTR+G+I substitution model for the concatenated data matrix, with four gamma categories, an uncorrelated relaxed-clock model and an incomplete sampling under birth-death models prior were applied to the specific Bayesian starting tree used as the tree prior. Four independent runs of 50 × 10^6^ generations each were performed, sampling every 1000th generation. Optimal convergence between four runs and the amount of burn-in were verified in Tracer v.1.6.0 (Drummond *et al.*, 2012) by checking effective sample size (ESS) scores and consistency of the results between multiple runs. We combined the four runs after discarding the initial 10% as burn-in, using Logcombiner 1.8.4 (Drummond *et al.*, 2012), and calculated a maximum clade-credibility tree with a posterior probability limit of 0.5, using TreeAnnotator v.1.8.4 (Drummond *et al.*, 2012), and visualised the tree using FigTree v.1.3.1(Rambaut & Drummond, 2010).

*Ancestral Character States Reconstruction*

The Symmetric (SYM), All Rates Different (ARD), and Equal Rate (ER) models of discrete character evolution, implemented in the *fitDiscrete* function of the geiger package (Harmon et al., 2008.), were tested for pollination syndrome states to identify the best fitting model of character evolution using a ML approach. The ER recognised as the best fit model in all reconstructions based on a AICc and AICc Weight (Table S5). Stochastic character mapping (1000 iterations) with the empirical Bayes method on the optimal model was carried out with the function *make.simmap* (Phytools) (Revell, 2012) in order to validate ML estimations.

*Models of trait evolution*

The fitted models compared the estimated parameters among supported models to test whether the evolutionary trajectories of floral traits vary among species with generalist, bird, long proboscis fly, and wind syndromes. To select the best model, we first applied stochastic character mapping (Huelsenbeck *et al.*, 2003) in an equal rates model (i.e. ER) to sample possible histories of the pollination syndromes on our phylogenetic tree (see above). Five models of BM_1_ (single-rate Brownian motion), BM_S_ (Brownian motion with different rate parameters for each state on a tree), OU_1_ (Ornstein-Uhlenbeck model with a single optimum for all species), OU_M_ (Ornstein-Uhlenbeck model with different state means and a single α and σ^2^ acting all selective regimes), and OU_MV_ (Ornstein-Uhlenbeck models that assume different state means as well as either multiple σ^2^) were considered in order to determine which aspects of the evolutionary trajectories of floral traits differ between pollination syndromes. OU_MA_ and OU_MVA_ models were considered too complex for the information contained in the data and were abandoned. We included the diagnostics of the eigendecomposition of the Hessian matrix in the runs to test if the information contained within the dataset is sufficient and parameters were estimated correctly. If the eigenvalues of the Hessian matrix are negative, parameter estimates are considered unreliable and the maximum likelihood estimate has not been found (Beaulieu *et al.*, 2012). All analyses were implemented using the R package OUwie (Beaulieu *et al.*, 2012).

**References**

**Beaulieu JM, Jhwueng DC, Boettiger C, O’Meara BC. 2012.** Modeling stabilizing selection: expanding the Ornstein–Uhlenbeck model of adaptive evolution. *Evolution: International Journal of Organic Evolution* **66**(8): 2369-2383.

**Breiman L. 2003.** Manual–setting up, using, and understanding random forests V4. 0. 2003 <http://oz>. berkeley. edu/users/breiman. *Using_random_forests_v4. 0. pdf*.

**Drummond AJ, Suchard MA, Xie D, Rambaut A. 2012.** Bayesian phylogenetics with BEAUti and the BEAST 1.7. *Molecular biology and evolution* **29**(8): 1969-1973.

**Giraudoux P 2017**. pgirmess: Data Analysis in Ecology. R package version 1.6.7.

**Huelsenbeck JP, Nielsen R, Bollback JP. 2003.** Stochastic mapping of morphological characters. *Systematic Biology* **52**(2): 131-158.

**Jung S, Marron JS. 2009.** PCA consistency in high dimension, low sample size context. *The Annals of Statistics* **37**(6B): 4104-4130.

**Klingenberg CP, Marugan-Lobon J. 2013.** Evolutionary Covariation in Geometric Morphometric Data: Analyzing Integration, Modularity, and Allometry in a Phylogenetic Context. *Systematic Biology* **62**(4): 591-610.

**Liaw A, Wiener M. 2002.** Classification and regression by randomForest. *R news* **2**(3): 18-22.

**Pirie M, Oliver E, De Kuppler AM, Gehrke B, Le Maitre N, Kandziora M, Bellstedt D. 2016.** The biodiversity hotspot as evolutionary hot-bed: spectacular radiation of Erica in the Cape Floristic Region. *BMC evolutionary biology* **16**(1): 190.

**R_Core_Team 2018**. R: A language and environment for statistical computing. . Vienna, Austria: R Foundation for Statistical Computing.

**Rambaut A, Drummond A. 2010.** FigTree v1. 3.1 Institute of Evolutionary Biology. *University of Edinburgh*.

**Revell LJ. 2012.** phytools: an R package for phylogenetic comparative biology (and other things). *Methods in Ecology and Evolution* **3**(2): 217-223.

**Staedler YM, Masson D, Schönenberger J. 2013.** Plant tissues in 3D via X-ray tomography: simple contrasting methods allow high resolution imaging. *PloS one* **8**(9): e75295.

**Young NM. 2006.** Function, ontogeny and canalization of shape variance in the primate scapula. *Journal of Anatomy* **209**(5): 623-636.
