## Supplementary tables (S1-S10) for "Modularity and evolution of flower shape: the role of efficiency, development, and spandrels in *Erica*"

**Table S1.** Species, sample numbers (n) and scanning conditions of *Erica* flowers. LFOV, large field of view objective. Voltage and current parameters refer to X-Ray source acceleration voltage and current, respectively.

| **sample** | **sample** | **species** | **n** | **Voltage [kV]** | **Current [µA]** | **Exposure time [s]** | **Pictures /sample** | **Pixel size [µm]** | **Objective** | **Binning** |
| --- | --- | --- | --- | --- | --- | --- | --- | --- | --- | --- |
| batch1 | A + B | *E. brachialis* | 8 | 40 | 200 | 6 | 728 | 15.5 | LFOV | 1 |
| batch2 | A | *E. curviflora* | 4 | 40 | 200 | 5 | 728 | 17.7 | LFOV | 1 |
| batch2 | B | *E. curviflora* | 3 | 40 | 200 | 5 | 728 | 18.4 | LFOV | 1 |
| batch3 | C | *E. perspicua* | 4 | 40 | 200 | 5 | 728 | 13.1 | LFOV | 1 |
| batch3 | B | *E. curvirostris* | 6 | 40 | 200 | 5 | 728 | 11.0 | LFOV | 1 |
| batch3 | A | *E. leucotrachela* | 4 | 40 | 200 | 5 | 728 | 14.9 | LFOV | 1 |
| batch4 | C | *E. curvirostris* | 6 | 40 | 200 | 5 | 728 | 17.4 | LFOV | 1 |
| batch4 | B | *E. curviflora* | 4 | 40 | 200 | 5 | 728 | 19.0 | LFOV | 1 |
| batch4 | A | *E. brachialis* | 4 | 40 | 200 | 6 | 728 | 17.4 | LFOV | 1 |
| batch5 | D+ E | *E. melanthera* | 13 | 40 | 200 | 5 | 728 | 11.3 | LFOV | 1 |
| batch5 | C | *E. blandfordia* | 5 | 40 | 200 | 5 | 728 | 11.3 | LFOV | 1 |
| batch5 | B | *E. hirtiflora* | 7 | 40 | 200 | 5 | 728 | 9.1 | LFOV | 1 |
| batch5 | A | *E. lateralis* | 6 | 40 | 200 | 5 | 728 | 11.3 | LFOV | 1 |
| batch6 | E | *E. turgida* | 8 | 40 | 200 | 5 | 728 | 10.3 | LFOV | 1 |
| batch6 | D | *E. lateralis* | 5 | 40 | 200 | 5 | 728 | 10.3 | LFOV | 1 |
| batch6 | C | *E. turgida* | 7 | 40 | 200 | 5 | 728 | 9.1 | LFOV | 1 |
| batch6 | B | *E. blandfordia* | 5 | 40 | 200 | 5 | 728 | 10.5 | LFOV | 1 |
| batch6 | A | *E. blandfordia* | 4 | 40 | 200 | 5 | 728 | 11.5 | LFOV | 1 |
| batch7 | A + B | *E. leucotrachela* | 8 | 40 | 200 | 5 | 728 | 17.8 | LFOV | 1 |
| batch8 | C | *E. perspicua* | 4 | 40 | 200 | 6 | 728 | 12.7 | LFOV | 1 |
| batch8 | A + B | *E. ventricosa* | 8 | 40 | 200 | 5 | 728 | 12.7 | LFOV | 1 |
| batch9 | D+E | *E. gracilis* | 12 | 40 | 200 | 6 | 728 | 10.5 | LFOV | 1 |
| batch9 | C | *E. hirtiflora* | 7 | 40 | 200 | 6 | 728 | 10.5 | LFOV | 1 |
| batch9 | B | *E. capensis* | 6 | 40 | 200 | 6 | 728 | 10.5 | LFOV | 1 |
| batch9 | A | *E. capensis* | 6 | 40 | 200 | 6 | 728 | 9.9 | LFOV | 1 |
| batch11 | C | *E. ventricosa* | 4 | 40 | 200 | 6 | 728 | 12.7 | LFOV | 1 |
| batch11 | B | *E. perspicua* | 4 | 40 | 200 | 6 | 728 | 13.3 | LFOV | 1 |
| batch12 | B | *E. gracilis* | 6 | 40 | 200 | 6 | 728 | 9.5 | 1x | 1 |
| batch12 | A | *E. gracilis* | 6 | 40 | 200 | 12 | 728 | 10.6 | 1x | 1 |
| batch13 | C | *E. spiculifolia* | 8 | 40 | 200 | 6 | 728 | 7.0 | LFOV | 1 |
| batch13 | B | *E. australis* | 4 | 40 | 200 | 4 | 728 | 10.9 | LFOV | 1 |
| batch13 | A | *E. australis* | 8 | 40 | 200 | 3 | 728 | 13.1 | LFOV | 1 |
| batch14 | E | *E. spiculifolia* | 8 | 40 | 200 | 15 | 728 | 9.7 | LFOV | 1 |
| batch14 | D | *E. scoparia* | 5 | 40 | 200 | 15 | 728 | 9.7 | LFOV | 1 |
| batch14 | A+B+C | *E. georgica* | 15 | 40 | 200 | 5 | 728 | 9.7 | LFOV | 1 |
| batch15 | E | *E. scoparia* | 8 | 40 | 200 | 5 | 728 | 9.2 | LFOV | 1 |
| batch15 | D | *E. vagans* | 7 | 40 | 200 | 5 | 728 | 8.8 | LFOV | 1 |
| batch15 | C | *E. vagans* | 7 | 40 | 200 | 5 | 728 | 8.7 | LFOV | 1 |
| batch15 | B | *E. melanthera* | 6 | 40 | 200 | 5 | 728 | 8.4 | LFOV | 1 |
| batch15 | A | *E. melanthera* | 7 | 40 | 200 | 5 | 728 | 8.8 | LFOV | 1 |

**Table S2.** Landmarks used to digitise the shape of *Erica* flowers (number, flower part, and position in frontal view), and modules to which they belong in the modularity hypotheses tested.

| **landmark** | **flower** | **position in** | **attraction/** | **developmental** | **efficiency 1** | **efficiency 2** |
| --- | --- | --- | --- | --- | --- | --- |
| **number** | **part** | **frontal view** | **reproduction** |  |  |  |
| 1 | base of flower |  | reproduction | carpels | remainder | carpels |
| 2 | nectary | upper | reproduction | carpels | remainder | carpels |
| 3 | nectary | lower | reproduction | carpels | remainder | carpels |
| 4 | ovary/style transition |  | reproduction | carpels | remainder | carpels |
| 5 | sepal tip | upper | attraction | sepals | remainder | remainder |
| 6 | sepal tip | left | attraction | sepals | remainder | remainder |
| 7 | sepal tip | lower | attraction | sepals | remainder | remainder |
| 8 | sepal tip | right | attraction | sepals | remainder | remainder |
| 9 | corolla - widest position | upper/left | attraction | petals | remainder | remainder |
| 10 | corolla - widest position | lower/left | attraction | petals | remainder | remainder |
| 11 | corolla - widest position | lower/right | attraction | petals | remainder | remainder |
| 12 | corolla - widest position | upper/right | attraction | petals | remainder | remainder |
| 13 | corolla - narrowest position | upper | attraction | petals | remainder | aperture |
| 14 | corolla - narrowest position | left | attraction | petals | remainder | aperture |
| 15 | corolla - narrowest position | lower | attraction | petals | remainder | aperture |
| 16 | corolla - narrowest position | right | attraction | petals | remainder | aperture |
| 17 | meeting of petal lobes | upper | attraction | petals | deposition | aperture |
| 18 | meeting of petal lobes | left | attraction | petals | deposition | aperture |
| 19 | meeting of petal lobes | lower | attraction | petals | receipt | aperture |
| 20 | meeting of petal lobes | right | attraction | petals | deposition | aperture |
| 21 | petal tip | upper/left | attraction | stamens | deposition | aperture |
| 22 | petal tip | lower/left | attraction | stamens | deposition | aperture |
| 23 | petal tip | lower/right | attraction | stamens | deposition | aperture |
| 24 | petal tip | upper/right | attraction | stamens | deposition | aperture |
| 25 | anther tip | upper | reproduction | stamens | deposition | remainder |
| 26 | anther tip | left | reproduction | stamens | deposition | remainder |
| 27 | anther tip | lower | reproduction | stamens | deposition | remainder |
| 28 | anther tip | right | reproduction | stamens | deposition | remainder |
| 29 | anther base | upper | reproduction | stamens | deposition | remainder |
| 30 | anther base | left | reproduction | stamens | deposition | remainder |
| 31 | anther base | lower | reproduction | stamens | deposition | remainder |
| 32 | anther base | right | reproduction | stamens | deposition | remainder |
| 33 | tip of stigma |  | reproduction | carpels | receipt | carpels |

**Table S3.** Species-level average values for size (centroid size) and integration (eigenvalue variance scaled by total variance and number of variables).

| Species | Centroid size | Integration |
| --- | --- | --- |
| Erica australis L. | 18,0 | 0,20 |
| Erica blandfordia Andrews | 16,2 | 0,17 |
| Erica bolusiae T. M. Salter | 8,9 | 0,24 |
| Erica brachialis Salisb. | 38,4 | 0,22 |
| Erica capensis T.M. Salter | 8,0 | 0,17 |
| Erica curviflora L. | 63,6 | 0,19 |
| Erica georgica L. Guthrie & Bolus | 18,3 | 0,14 |
| Erica gracilis J.C. Wendl. | 7,9 | 0,18 |
| Erica hirtiflora Curtis | 7,9 | 0,29 |
| Erica lateralis Willd. | 11,9 | 0,16 |
| Erica leucotrachela H.A. Baker | 42,0 | 0,21 |
| Erica margaritacea Aiton | 12,3 | 0,32 |
| Erica melanthera L. | 8,3 | 0,33 |
| Erica perspicua J.C. Wendl. | 44,0 | 0,16 |
| Erica scoparia L. | 4,4 | 0,24 |
| Erica spiculifolia Salisb. | 5,7 | 0,26 |
| Erica turgida Salisb. | 8,0 | 0,14 |
| Erica vagans L. | 8,7 | 0,27 |
| Erica ventricosa Thunb. | 29,8 | 0,23 |

**Table S4.** Genbank accession numbers for nrDNA ITS and cpDNA trnL-F-ndhJ and trnT-L sequence data. The dash symbol (–) shows that the sequence is not available.

| **Species** | **Source or Specimen** | **ITS** | ***trn*LF-ndhJ** | ***trnT-L*** |
| --- | --- | --- | --- | --- |
| *Calluna vulgaris* | Pirie et al., 2016 | HQ858882 | KP737378 | _ |
| *Daboecia cantabrica* | Pirie et al., 2016 | HQ859000 | KP737380 | KP737653 |
| *Erica australis* | Pirie et al., 2016 | HQ858926 | HQ858927 | HQ858928 |
| *Erica blandfordii* | Pirie et al., 2016 | HQ858940 | KU832634 | KU831854 |
| *Erica bolusiae* | Pirie et al., 2016 | KU832354 | KU832640 | KU831859 |
| *Erica brachialis* | Pirie et al., 2016 | HQ858944 | KU832642 | KU831861 |
| *Erica capensis* | Pirie et al., 2016 | KU832452 | KU832846 | KU832055 |
| *Erica curviflora* | Pirie et al., 2016 | KU832381 | KU832698 | KU831913 |
| *Erica georgica* | Pirie et al., 2016 | KU832410 | KU832764 | KU831977 |
| *Erica gracilis* | Pirie et al., 2016 | HQ859074 | KU832788 | KU832000 |
| *Erica hirtiflora* | Pirie et al., 2016 | HQ859084 | KU832808 | KU832019 |
| *Erica lateralis* | Pirie et al., 2016 | KU832456 | KU832851 | KU832060 |
| *Erica leucotrachela* | Pirie et al., 2016 | KU832461 | KU832865 | _ |
| *Erica margaritacea* | Pirie et al., 2016 | KU832468 | KU832882 | KU832088 |
| *Erica melanthera* | Pirie et al., 2016 | HQ859145 | KU832889 | KU832094 |
| *Erica perspicua* | Pirie et al., 2016 | HQ859192 | KU832949 | KU832153 |
| *Erica scoparia* | Mugrabi de Kuppler et al., 2015 | KP737593 | KP737459 | KP737674 |
| *Erica spiculifolia* | Mugrabi de Kuppler et al., 2015 | KP737610 | KP737475 | KP737678 |
| *Erica turgida* | Pirie et al., 2016 | HQ859312 | KU833085 | KU832288 |
| *Erica vagans* | Pirie et al., 2016 | HQ859319 | KP737490 | KP737682 |
| *Erica ventricosa* | Pirie et al., 2016 | KU832562 | KU833104 | KU832307 |

**Table S5.** Discrete character mapping models for pollination syndromes. Model testing (equal-rates model – ER, symmetrical rates model – SYM and all-rates-different model – ARD) for ancestral state reconstruction under the ML approach. The ER model is preferred (lowest

AICc and AICc Weight values). AICc = corrected AIC score.

|  | AICc | AICc Weight |
| --- | --- | --- |
| **ER** | **39.1** | **0.99955** |
| SYM | 54.6 | 0.00045 |
| ARD | 108 | 0 |

**Table S6.** Main variables mean accuracy decrease of random forest syndrome prediction (averaged on 500 RF made of 1001 trees each).

| Variable | Mean (± SD) decrease |
| --- | --- |
|  | in accuracy |
| tube length | 0.0442 ± 0.0036 |
| z16 - corolla narrowest position | 0.0296 ± 0.0031 |
| z14 - corolla narrowest position | 0.0297 ± 0.0031 |
| z9 - corolla widest position | 0.0279 ± 0.0027 |
| z12 - corolla widest position | 0.0281 ± 0.0028 |
| y13 - corolla narrowest position | 0.0248 ± 0.0031 |
| x4 - ovary/style transition | 0.0248 ± 0.0025 |
| y15 - corolla narrowest position | 0.0234 ± 0.0030 |
| z20 - meeting of petal lobes | 0.0204 ± 0.0001 |
| z11 - corolla widest position | 0.0204 ± 0.0028 |
| z18 - meeting of petal lobes | 0.0206 ± 0.0024 |
| z10 - corolla widest position | 0.0206 ± 0.0023 |
| y7 - sepal tip | 0.0190 ± 0.0022 |
| y11 - corolla widest position | 0.0176 ± 0.0021 |
| y10 - corolla widest position | 0.0175 ± 0.0023 |
| y9 - corolla widest position | 0.0172 ± 0.0022 |

**Table S7.** Corolla tube length per flower calculated as the distance between the base of the flower and the average position of the meeting of the four corolla lobes.

| flower number | *Erica* species | tube length |
| --- | --- | --- |
| 1 | *australis* | 6.27 |
| 2 | *australis* | 6.92 |
| 3 | *australis* | 6.58 |
| 4 | *australis* | 6.99 |
| 5 | *australis* | 7.03 |
| 6 | *australis* | 7.10 |
| 7 | *australis* | 6.82 |
| 8 | *australis* | 6.57 |
| 9 | *australis* | 6.88 |
| 10 | *australis* | 6.85 |
| 11 | *australis* | 6.87 |
| 12 | *capensis* | 2.10 |
| 13 | *capensis* | 2.09 |
| 14 | *capensis* | 2.13 |
| 15 | *capensis* | 2.07 |
| 16 | *capensis* | 2.05 |
| 17 | *capensis* | 2.17 |
| 18 | *capensis* | 2.15 |
| 19 | *capensis* | 2.06 |
| 20 | *capensis* | 1.97 |
| 21 | *capensis* | 2.13 |
| 22 | *blandifordia* | 6.23 |
| 23 | *blandifordia* | 6.62 |
| 24 | *blandifordia* | 6.03 |
| 25 | *blandifordia* | 6.38 |
| 26 | *blandifordia* | 6.06 |
| 27 | *blandifordia* | 6.69 |
| 28 | *blandifordia* | 6.63 |
| 29 | *blandifordia* | 6.17 |
| 30 | *blandifordia* | 6.35 |
| 31 | *blandifordia* | 6.44 |
| 32 | *blandifordia* | 5.72 |
| 33 | *brachialis* | 18.21 |
| 34 | *brachialis* | 18.97 |
| 35 | *brachialis* | 18.07 |
| 36 | *brachialis* | 18.91 |
| 37 | *brachialis* | 18.19 |
| 38 | *brachialis* | 17.97 |
| 39 | *brachialis* | 19.21 |
| 40 | *brachialis* | 18.05 |
| 41 | *brachialis* | 19.13 |
| 42 | *brachialis* | 19.16 |
| 43 | *brachialis* | 18.02 |
| 44 | *brachialis* | 17.08 |
| 45 | *brachialis* | 17.97 |
| 46 | *brachialis* | 18.36 |
| 47 | *curviflora* | 21.71 |
| 48 | *curviflora* | 22.37 |
| 49 | *curviflora* | 21.30 |
| 50 | *curviflora* | 22.36 |
| 51 | *curviflora* | 22.78 |
| 52 | *curviflora* | 22.55 |
| 53 | *curviflora* | 22.93 |
| 54 | *curviflora* | 22.31 |
| 55 | *curviflora* | 20.49 |
| 56 | *curviflora* | 23.42 |
| 57 | *curviflora* | 20.86 |
| 58 | *bolusiae* | 3.43 |
| 59 | *bolusiae* | 3.54 |
| 60 | *bolusiae* | 3.56 |
| 61 | *bolusiae* | 3.31 |
| 62 | *bolusiae* | 3.37 |
| 63 | *bolusiae* | 3.16 |
| 64 | *bolusiae* | 3.37 |
| 65 | *bolusiae* | 3.29 |
| 66 | *bolusiae* | 2.89 |
| 67 | *bolusiae* | 3.07 |
| 68 | *georgica* | 9.43 |
| 69 | *georgica* | 8.75 |
| 70 | *georgica* | 9.25 |
| 71 | *georgica* | 8.62 |
| 72 | *georgica* | 9.37 |
| 73 | *georgica* | 8.59 |
| 74 | *georgica* | 8.43 |
| 75 | *georgica* | 8.70 |
| 76 | *georgica* | 9.13 |
| 77 | *georgica* | 9.35 |
| 78 | *georgica* | 7.84 |
| 79 | *georgica* | 8.57 |
| 80 | *georgica* | 8.48 |
| 81 | *georgica* | 9.02 |
| 82 | *georgica* | 8.84 |
| 85 | *gracilis* | 2.85 |
| 86 | *gracilis* | 3.26 |
| 87 | *gracilis* | 2.84 |
| 88 | *gracilis* | 2.78 |
| 89 | *gracilis* | 3.21 |
| 90 | *gracilis* | 3.22 |
| 91 | *gracilis* | 3.26 |
| 92 | *gracilis* | 3.08 |
| 93 | *gracilis* | 3.15 |
| 94 | *gracilis* | 3.25 |
| 95 | *hirtiflora* | 2.61 |
| 96 | *hirtiflora* | 2.97 |
| 97 | *hirtiflora* | 2.80 |
| 98 | *hirtiflora* | 2.82 |
| 99 | *hirtiflora* | 2.81 |
| 100 | *hirtiflora* | 2.92 |
| 101 | *hirtiflora* | 2.82 |
| 102 | *hirtiflora* | 2.78 |
| 103 | *hirtiflora* | 2.68 |
| 104 | *hirtiflora* | 2.54 |
| 105 | *lateralis* | 3.89 |
| 106 | *lateralis* | 3.56 |
| 107 | *lateralis* | 3.66 |
| 108 | *lateralis* | 3.97 |
| 109 | *lateralis* | 3.49 |
| 110 | *lateralis* | 3.76 |
| 111 | *lateralis* | 3.58 |
| 112 | *lateralis* | 3.68 |
| 113 | *lateralis* | 3.79 |
| 114 | *lateralis* | 3.54 |
| 115 | *leucotrachela* | 17.60 |
| 116 | *leucotrachela* | 18.41 |
| 117 | *leucotrachela* | 18.27 |
| 118 | *leucotrachela* | 17.52 |
| 119 | *leucotrachela* | 16.85 |
| 120 | *leucotrachela* | 17.82 |
| 121 | *leucotrachela* | 16.75 |
| 122 | *leucotrachela* | 18.06 |
| 123 | *leucotrachela* | 17.48 |
| 124 | *leucotrachela* | 19.73 |
| 125 | *melanthera* | 2.80 |
| 126 | *melanthera* | 2.72 |
| 127 | *melanthera* | 2.71 |
| 128 | *melanthera* | 2.69 |
| 129 | *melanthera* | 2.60 |
| 130 | *melanthera* | 2.91 |
| 131 | *melanthera* | 2.77 |
| 132 | *melanthera* | 2.87 |
| 133 | *melanthera* | 3.41 |
| 134 | *melanthera* | 2.77 |
| 135 | *perspicua* | 17.92 |
| 136 | *perspicua* | 18.70 |
| 137 | *perspicua* | 17.61 |
| 138 | *perspicua* | 18.57 |
| 139 | *perspicua* | 15.69 |
| 140 | *perspicua* | 17.29 |
| 141 | *perspicua* | 17.40 |
| 142 | *perspicua* | 18.62 |
| 143 | *perspicua* | 14.38 |
| 144 | *perspicua* | 17.61 |
| 145 | *margaritacea* | 2.90 |
| 146 | *margaritacea* | 2.91 |
| 147 | *margaritacea* | 4.18 |
| 148 | *margaritacea* | 4.15 |
| 149 | *margaritacea* | 3.74 |
| 150 | *margaritacea* | 2.88 |
| 151 | *margaritacea* | 3.99 |
| 152 | *margaritacea* | 4.08 |
| 153 | *margaritacea* | 2.80 |
| 154 | *margaritacea* | 4.12 |
| 155 | *margaritacea* | 3.60 |
| 156 | *margaritacea* | 2.72 |
| 157 | *margaritacea* | 2.84 |
| 158 | *scoparia* | 1.20 |
| 159 | *scoparia* | 1.29 |
| 160 | *scoparia* | 1.26 |
| 161 | *scoparia* | 1.27 |
| 162 | *scoparia* | 1.28 |
| 163 | *scoparia* | 1.26 |
| 164 | *scoparia* | 1.14 |
| 165 | *scoparia* | 1.29 |
| 166 | *scoparia* | 1.21 |
| 167 | *scoparia* | 1.15 |
| 168 | *spiculifolia* | 1.78 |
| 169 | *spiculifolia* | 1.66 |
| 170 | *spiculifolia* | 1.79 |
| 171 | *spiculifolia* | 1.77 |
| 172 | *spiculifolia* | 1.76 |
| 173 | *spiculifolia* | 1.76 |
| 174 | *spiculifolia* | 1.87 |
| 175 | *spiculifolia* | 1.73 |
| 176 | *spiculifolia* | 1.78 |
| 177 | *spiculifolia* | 1.74 |
| 178 | *spiculifolia* | 1.81 |
| 179 | *spiculifolia* | 1.83 |
| 180 | *turgida* | 2.30 |
| 181 | *turgida* | 2.31 |
| 182 | *turgida* | 2.11 |
| 183 | *turgida* | 2.03 |
| 184 | *turgida* | 2.25 |
| 185 | *turgida* | 2.03 |
| 186 | *turgida* | 2.21 |
| 187 | *turgida* | 2.20 |
| 188 | *turgida* | 2.18 |
| 189 | *turgida* | 2.09 |
| 190 | *turgida* | 2.14 |
| 191 | *turgida* | 2.25 |
| 192 | *vagans* | 1.90 |
| 193 | *vagans* | 2.01 |
| 194 | *vagans* | 1.99 |
| 195 | *vagans* | 2.17 |
| 196 | *vagans* | 1.96 |
| 197 | *vagans* | 1.98 |
| 198 | *vagans* | 1.99 |
| 199 | *vagans* | 1.82 |
| 200 | *vagans* | 1.86 |
| 201 | *vagans* | 1.97 |
| 202 | *vagans* | 1.99 |
| 204 | *ventricosa* | 13.83 |
| 205 | *ventricosa* | 13.93 |
| 206 | *ventricosa* | 13.78 |
| 207 | *ventricosa* | 13.74 |
| 208 | *ventricosa* | 13.94 |
| 209 | *ventricosa* | 13.73 |
| 210 | *ventricosa* | 14.27 |
| 211 | *ventricosa* | 13.83 |
| 212 | *ventricosa* | 13.46 |

**Table S8.** Classification of 114 individual flowers of diverse *Erica* species into the pollination syndromes, based on floral shape and size; proportions of assignation to each of the syndromes for each individual calculated based on predictions from 500 random forests of 1001 trees each.

| flower number | *Erica* species | generalist | bird | long-proboscid fly | wind | sum |
| --- | --- | --- | --- | --- | --- | --- |
| 22 | *blandifordia* | 1 | 0 | 0 | 0 | 1 |
| 23 | *blandifordia* | 1 | 0 | 0 | 0 | 1 |
| 24 | *blandifordia* | 1 | 0 | 0 | 0 | 1 |
| 25 | *blandifordia* | 1 | 0 | 0 | 0 | 1 |
| 26 | *blandifordia* | 1 | 0 | 0 | 0 | 1 |
| 27 | *blandifordia* | 1 | 0 | 0 | 0 | 1 |
| 28 | *blandifordia* | 1 | 0 | 0 | 0 | 1 |
| 29 | *blandifordia* | 1 | 0 | 0 | 0 | 1 |
| 30 | *blandifordia* | 0 | 0 | 1 | 0 | 1 |
| 31 | *blandifordia* | 1 | 0 | 0 | 0 | 1 |
| 32 | *blandifordia* | 1 | 0 | 0 | 0 | 1 |
| 47 | *curviflora* | 0 | 1 | 0 | 0 | 1 |
| 48 | *curviflora* | 0 | 1 | 0 | 0 | 1 |
| 49 | *curviflora* | 0 | 1 | 0 | 0 | 1 |
| 50 | *curviflora* | 0 | 1 | 0 | 0 | 1 |
| 51 | *curviflora* | 0 | 1 | 0 | 0 | 1 |
| 52 | *curviflora* | 0 | 1 | 0 | 0 | 1 |
| 53 | *curviflora* | 0 | 1 | 0 | 0 | 1 |
| 54 | *curviflora* | 0 | 1 | 0 | 0 | 1 |
| 55 | *curviflora* | 0 | 1 | 0 | 0 | 1 |
| 56 | *curviflora* | 0 | 1 | 0 | 0 | 1 |
| 57 | *curviflora* | 0 | 1 | 0 | 0 | 1 |
| 58 | *bolusiae* | 1 | 0 | 0 | 0 | 1 |
| 59 | *bolusiae* | 1 | 0 | 0 | 0 | 1 |
| 60 | *bolusiae* | 1 | 0 | 0 | 0 | 1 |
| 61 | *bolusiae* | 1 | 0 | 0 | 0 | 1 |
| 62 | *bolusiae* | 1 | 0 | 0 | 0 | 1 |
| 63 | *bolusiae* | 1 | 0 | 0 | 0 | 1 |
| 64 | *bolusiae* | 1 | 0 | 0 | 0 | 1 |
| 65 | *bolusiae* | 1 | 0 | 0 | 0 | 1 |
| 66 | *bolusiae* | 1 | 0 | 0 | 0 | 1 |
| 67 | *bolusiae* | 1 | 0 | 0 | 0 | 1 |
| 68 | *georgica* | 0 | 1 | 0 | 0 | 1 |
| 69 | *georgica* | 0.016 | 0.294 | 0.69 | 0 | 1 |
| 70 | *georgica* | 0.007 | 0.993 | 0 | 0 | 1 |
| 71 | *georgica* | 0.061 | 0.939 | 0 | 0 | 1 |
| 72 | *georgica* | 0.049 | 0.951 | 0 | 0 | 1 |
| 73 | *georgica* | 0.239 | 0.759 | 0.002 | 0 | 1 |
| 74 | *georgica* | 0 | 1 | 0 | 0 | 1 |
| 75 | *georgica* | 0.338 | 0.662 | 0 | 0 | 1 |
| 76 | *georgica* | 0 | 1 | 0 | 0 | 1 |
| 77 | *georgica* | 0 | 1 | 0 | 0 | 1 |
| 78 | *georgica* | 1 | 0 | 0 | 0 | 1 |
| 79 | *georgica* | 0.994 | 0.006 | 0 | 0 | 1 |
| 80 | *georgica* | 1 | 0 | 0 | 0 | 1 |
| 81 | *georgica* | 0 | 1 | 0 | 0 | 1 |
| 82 | *georgica* | 0.018 | 0.982 | 0 | 0 | 1 |
| 105 | *lateralis* | 1 | 0 | 0 | 0 | 1 |
| 106 | *lateralis* | 1 | 0 | 0 | 0 | 1 |
| 107 | *lateralis* | 1 | 0 | 0 | 0 | 1 |
| 108 | *lateralis* | 1 | 0 | 0 | 0 | 1 |
| 109 | *lateralis* | 1 | 0 | 0 | 0 | 1 |
| 110 | *lateralis* | 1 | 0 | 0 | 0 | 1 |
| 111 | *lateralis* | 1 | 0 | 0 | 0 | 1 |
| 112 | *lateralis* | 1 | 0 | 0 | 0 | 1 |
| 113 | *lateralis* | 1 | 0 | 0 | 0 | 1 |
| 114 | *lateralis* | 1 | 0 | 0 | 0 | 1 |
| 115 | *leucotrachela* | 0 | 1 | 0 | 0 | 1 |
| 116 | *leucotrachela* | 0 | 1 | 0 | 0 | 1 |
| 117 | *leucotrachela* | 0 | 1 | 0 | 0 | 1 |
| 118 | *leucotrachela* | 0 | 1 | 0 | 0 | 1 |
| 119 | *leucotrachela* | 0 | 1 | 0 | 0 | 1 |
| 120 | *leucotrachela* | 0 | 1 | 0 | 0 | 1 |
| 121 | *leucotrachela* | 0 | 1 | 0 | 0 | 1 |
| 122 | *leucotrachela* | 0 | 1 | 0 | 0 | 1 |
| 123 | *leucotrachela* | 0 | 1 | 0 | 0 | 1 |
| 124 | *leucotrachela* | 0 | 1 | 0 | 0 | 1 |
| 125 | *melanthera* | 1 | 0 | 0 | 0 | 1 |
| 126 | *melanthera* | 1 | 0 | 0 | 0 | 1 |
| 127 | *melanthera* | 1 | 0 | 0 | 0 | 1 |
| 128 | *melanthera* | 1 | 0 | 0 | 0 | 1 |
| 129 | *melanthera* | 1 | 0 | 0 | 0 | 1 |
| 130 | *melanthera* | 1 | 0 | 0 | 0 | 1 |
| 131 | *melanthera* | 1 | 0 | 0 | 0 | 1 |
| 132 | *melanthera* | 1 | 0 | 0 | 0 | 1 |
| 133 | *melanthera* | 1 | 0 | 0 | 0 | 1 |
| 134 | *melanthera* | 1 | 0 | 0 | 0 | 1 |
| 145 | *margaritacea* | 1 | 0 | 0 | 0 | 1 |
| 146 | *margaritacea* | 1 | 0 | 0 | 0 | 1 |
| 147 | *margaritacea* | 1 | 0 | 0 | 0 | 1 |
| 148 | *margaritacea* | 1 | 0 | 0 | 0 | 1 |
| 149 | *margaritacea* | 1 | 0 | 0 | 0 | 1 |
| 150 | *margaritacea* | 1 | 0 | 0 | 0 | 1 |
| 151 | *margaritacea* | 1 | 0 | 0 | 0 | 1 |
| 152 | *margaritacea* | 1 | 0 | 0 | 0 | 1 |
| 153 | *margaritacea* | 1 | 0 | 0 | 0 | 1 |
| 154 | *margaritacea* | 1 | 0 | 0 | 0 | 1 |
| 155 | *margaritacea* | 1 | 0 | 0 | 0 | 1 |
| 156 | *margaritacea* | 1 | 0 | 0 | 0 | 1 |
| 157 | *margaritacea* | 1 | 0 | 0 | 0 | 1 |
| 168 | *spiculifolia* | 1 | 0 | 0 | 0 | 1 |
| 169 | *spiculifolia* | 1 | 0 | 0 | 0 | 1 |
| 170 | *spiculifolia* | 1 | 0 | 0 | 0 | 1 |
| 171 | *spiculifolia* | 1 | 0 | 0 | 0 | 1 |
| 172 | *spiculifolia* | 1 | 0 | 0 | 0 | 1 |
| 173 | *spiculifolia* | 1 | 0 | 0 | 0 | 1 |
| 174 | *spiculifolia* | 1 | 0 | 0 | 0 | 1 |
| 175 | *spiculifolia* | 1 | 0 | 0 | 0 | 1 |
| 176 | *spiculifolia* | 1 | 0 | 0 | 0 | 1 |
| 177 | *spiculifolia* | 1 | 0 | 0 | 0 | 1 |
| 178 | *spiculifolia* | 1 | 0 | 0 | 0 | 1 |
| 179 | *spiculifolia* | 1 | 0 | 0 | 0 | 1 |
| 180 | *turgida* | 1 | 0 | 0 | 0 | 1 |
| 181 | *turgida* | 1 | 0 | 0 | 0 | 1 |
| 182 | *turgida* | 1 | 0 | 0 | 0 | 1 |
| 183 | *turgida* | 1 | 0 | 0 | 0 | 1 |
| 184 | *turgida* | 1 | 0 | 0 | 0 | 1 |
| 185 | *turgida* | 1 | 0 | 0 | 0 | 1 |
| 186 | *turgida* | 1 | 0 | 0 | 0 | 1 |
| 187 | *turgida* | 1 | 0 | 0 | 0 | 1 |
| 188 | *turgida* | 1 | 0 | 0 | 0 | 1 |
| 189 | *turgida* | 1 | 0 | 0 | 0 | 1 |
| 190 | *turgida* | 1 | 0 | 0 | 0 | 1 |
| 191 | *turgida* | 1 | 0 | 0 | 0 | 1 |

**Table S9.** Support values evolutionary models of floral shape evolution. Support for BM, OU, and EB models for the evolution of highly-dimensional whole floral shape in *Erica*; preferred model (lowest GIC value) in bold.).

|  | GIC | Log-likelihood | parameters |
| --- | --- | --- | --- |
| BM | -13276 | 6766 | - |
| **OU** | **-13325** | **6793** | **Alpha = 2.57** |
| EB | -13274 | 6766 | r = 0 |

**Table S10.** Summary of the preferred models of evolution for seven phenotypic trait variables (PC1-5 of floral shape, centroid size, and integration) under the pollination-syndrome regime. Parameter estimates are reported as a mean ± standard error. Models of quantitative trait evolution with their parameters, indicating for each model whether  (the optimum trait value), σ² (the intensity of random fluctuation in the evolutionary trajectory), and α (the selective pull toward the optimal value) are modelled with one global parameter (BM_1_ and OU_1_) or four parameters (OU_M_) that are specific for each pollination syndrome.

| Shape and size | Model | θ | | | | | σ² | α | |
| --- | --- | --- | --- | --- | --- | --- | --- | --- | --- |
| variables |  | Generalist | Bird | LPF | Wind | |  |  |  |
| Centroid size | OUM | 10,36±1,56 | 49,95±2,91 | 31,92±6,63 | -2,51±7,13 | 95.79 | | | 1.63 |
| PC1 (39% variance) | OUM | 0,09±0,03 | -0,35±0,06 | -0,72±0,25 | 0,50±0,15 | 1.36E-02 | | | 0.68 |
| PC2 (22% variance) | OU1 | 3.49E-03±3.47E-02 | | | | | 1.23E-02 | 0.32 | |
| PC3 (11% variance) | BM1 | 5.35E-02±6.19E-02 | | | | | 7.54E-04 | - | |
| PC4 (7% variance) | BM1 | 2.85E-02±5.66E-02 | | | | | 6.32E-04 | - | |
| PC5 (6% variance) | OU1 | 1.41E-03±1.78E-02 | | | | | 3.71E-03 | 0.35 | |
| Integration | OU1 | 0,23±0,02 | | | | | 1.15E-03 | 0.18 | |
