## Supplementary material for "Modularity and evolution of flower shape: the role of efficiency, development, and spandrels in *Erica*": Notes S1 (supplementary data)

**Notes S1: literature analysis**

Search in ISI Web of Knowledge for keyword “geometric morphometrics”, refined by year and topic (“zoology”, “anthropology”, and “plant sciences”)

| year | zoology | anthropology | plant sciences |
| --- | --- | --- | --- |
| 1990 | 1 | 0 | 0 |
| 1991 | 2 | 0 | 0 |
| 1992 | 0 | 0 | 0 |
| 1993 | 0 | 1 | 0 |
| 1994 | 0 | 0 | 0 |
| 1995 | 0 | 1 | 0 |
| 1996 | 8 | 2 | 0 |
| 1997 | 1 | 0 | 0 |
| 1998 | 9 | 0 | 0 |
| 1999 | 4 | 1 | 0 |
| 2000 | 6 | 5 | 0 |
| 2001 | 7 | 2 | 0 |
| 2002 | 6 | 7 | 1 |
| 2003 | 6 | 8 | 1 |
| 2004 | 21 | 10 | 0 |
| 2005 | 21 | 12 | 3 |
| 2006 | 13 | 10 | 2 |
| 2007 | 14 | 22 | 5 |
| 2008 | 29 | 22 | 2 |
| 2009 | 36 | 19 | 13 |
| 2010 | 32 | 27 | 12 |
| 2011 | 45 | 31 | 4 |
| 2012 | 55 | 29 | 8 |
| 2013 | 64 | 45 | 12 |
| 2014 | 48 | 35 | 7 |
| 2015 | 72 | 48 | 5 |
| 2016 | 70 | 46 | 13 |
| 2017 | 82 | 46 | 4 |
| 2018 | 70 | 41 | 14 |

Keyword "geometric morphometrics" refined by topic

|  | zoology | anthropology | plant sciences |
| --- | --- | --- | --- |
| 1990 | 1 | 0 | 0 |
| 1991 | 2 | 0 | 0 |
| 1992 | 0 | 0 | 0 |
| 1993 | 0 | 1 | 0 |
| 1994 | 0 | 0 | 0 |
| 1995 | 0 | 1 | 0 |
| 1996 | 8 | 2 | 0 |
| 1997 | 1 | 0 | 0 |
| 1998 | 9 | 0 | 0 |
| 1999 | 4 | 1 | 0 |
| 2000 | 6 | 5 | 0 |
| 2001 | 7 | 2 | 0 |
| 2002 | 6 | 7 | 1 |
| 2003 | 6 | 8 | 1 |
| 2004 | 21 | 10 | 0 |
| 2005 | 21 | 12 | 3 |
| 2006 | 13 | 10 | 2 |
| 2007 | 14 | 22 | 5 |
| 2008 | 29 | 22 | 2 |
| 2009 | 36 | 19 | 13 |
| 2010 | 32 | 27 | 12 |
| 2011 | 45 | 31 | 4 |
| 2012 | 55 | 29 | 8 |
| 2013 | 64 | 45 | 12 |
| 2014 | 48 | 35 | 7 |
| 2015 | 72 | 48 | 5 |
| 2016 | 70 | 46 | 13 |
| 2017 | 82 | 46 | 4 |
| 2018 | 70 | 41 | 14 |

Search in ISI Web of Knowledge for keyword “modularity” and “integration” refined by year and topic (“zoology”, “anthropology”, and “plant sciences”)

| year | zoology | anthropology | plant sciences |
| --- | --- | --- | --- |
| 1990 | 0 | 0 | 0 |
| 1991 | 0 | 0 | 0 |
| 1992 | 0 | 0 | 0 |
| 1993 | 0 | 0 | 0 |
| 1994 | 0 | 0 | 0 |
| 1995 | 0 | 0 | 0 |
| 1996 | 0 | 0 | 0 |
| 1997 | 0 | 0 | 0 |
| 1998 | 0 | 0 | 0 |
| 1999 | 0 | 0 | 0 |
| 2000 | 1 | 0 | 0 |
| 2001 | 1 | 0 | 0 |
| 2002 | 1 | 1 | 0 |
| 2003 | 2 | 0 | 0 |
| 2004 | 6 | 0 | 0 |
| 2005 | 0 | 2 | 1 |
| 2006 | 0 | 1 | 0 |
| 2007 | 3 | 0 | 1 |
| 2008 | 3 | 4 | 1 |
| 2009 | 5 | 4 | 1 |
| 2010 | 5 | 3 | 1 |
| 2011 | 5 | 7 | 0 |
| 2012 | 7 | 3 | 1 |
| 2013 | 4 | 7 | 2 |
| 2014 | 7 | 5 | 0 |
| 2015 | 9 | 4 | 0 |
| 2016 | 5 | 1 | 6 |
| 2017 | 5 | 3 | 0 |
| 2018 | 5 | 8 | 1 |

Keywords "modularity" & "integration"

|  | zoology | anthropology | plant sciences |
| --- | --- | --- | --- |
| 1990 | 0 | 0 | 0 |
| 1991 | 0 | 0 | 0 |
| 1992 | 0 | 0 | 0 |
| 1993 | 0 | 0 | 0 |
| 1994 | 0 | 0 | 0 |
| 1995 | 0 | 0 | 0 |
| 1996 | 0 | 0 | 0 |
| 1997 | 0 | 0 | 0 |
| 1998 | 0 | 0 | 0 |
| 1999 | 0 | 0 | 0 |
| 2000 | 1 | 0 | 0 |
| 2001 | 1 | 0 | 0 |
| 2002 | 1 | 1 | 0 |
| 2003 | 2 | 0 | 0 |
| 2004 | 6 | 0 | 0 |
| 2005 | 0 | 2 | 1 |
| 2006 | 0 | 1 | 0 |
| 2007 | 3 | 0 | 1 |
| 2008 | 3 | 4 | 1 |
| 2009 | 5 | 4 | 1 |
| 2010 | 5 | 3 | 1 |
| 2011 | 5 | 7 | 0 |
| 2012 | 7 | 3 | 1 |
| 2013 | 4 | 7 | 2 |
| 2014 | 7 | 5 | 0 |
| 2015 | 9 | 4 | 0 |
| 2016 | 5 | 1 | 6 |
| 2017 | 5 | 3 | 0 |
| 2018 | 5 | 8 | 1 |
