## Supplementary material for "Modularity and evolution of flower shape: the role of efficiency, development, and spandrels in *Erica*": Notes S1 (supplementary data)

**Notes S2 Allometric regressions and correlation between the corolla tube length and centroid size**

**Allometric regressions**

The proportion of variation explained by allometry (pooled by species) differs according to syndrome: bees: 1.53% (*P* = 0.01); birds: 2.96% (*P* = 0.19); long proboscis flies: 11.92% (*P* = 0.00032); wind: 23.23% (*P* = 0.047), details below:

**Only species with bee syndrome, variation pooled by species**

Sums of squares (the sums of squares below are within-group SSs)

Total SS:	1.81339384

Predicted SS:	0.02778468

Residual SS:	1.78560916

% predicted:	1.5322%

Permutation test against the null hypothesis of independence

Number of randomization rounds: 100000

P-value: 0.00968

**Only species with bird syndrome, variation pooled by species**

Sums of squares (the sums of squares below are within-group SSs)

Total SS:	0.27116450

Predicted SS:	0.00806889

Residual SS:	0.26309561

% predicted:	2.9756%

Permutation test against the null hypothesis of independence

Number of randomization rounds: 100000

P-value: 0.18973

**Only species with long proboscis fly syndrome, variation pooled by species**

Sums of squares (the sums of squares below are within-group SSs)

Total SS:	0.20556076

Predicted SS:	0.02448754

Residual SS:	0.18107322

% predicted:	11.9126%

Permutation test against the null hypothesis of independence

Number of randomization rounds: 100000

P-value: 0.00036

**Only species with wind syndrome, variation pooled by species**

Sums of squares (the sums of squares below are within-group SSs)

Total SS:	0.08374433

Predicted SS:	0.01945447

Residual SS:	0.06428987

% predicted:	23.2308%

Permutation test against the null hypothesis of independence

Number of randomization rounds: 100000

P-value: 0.04580

**Correlation between the corolla tube length and centroid size for all studied flowers together**

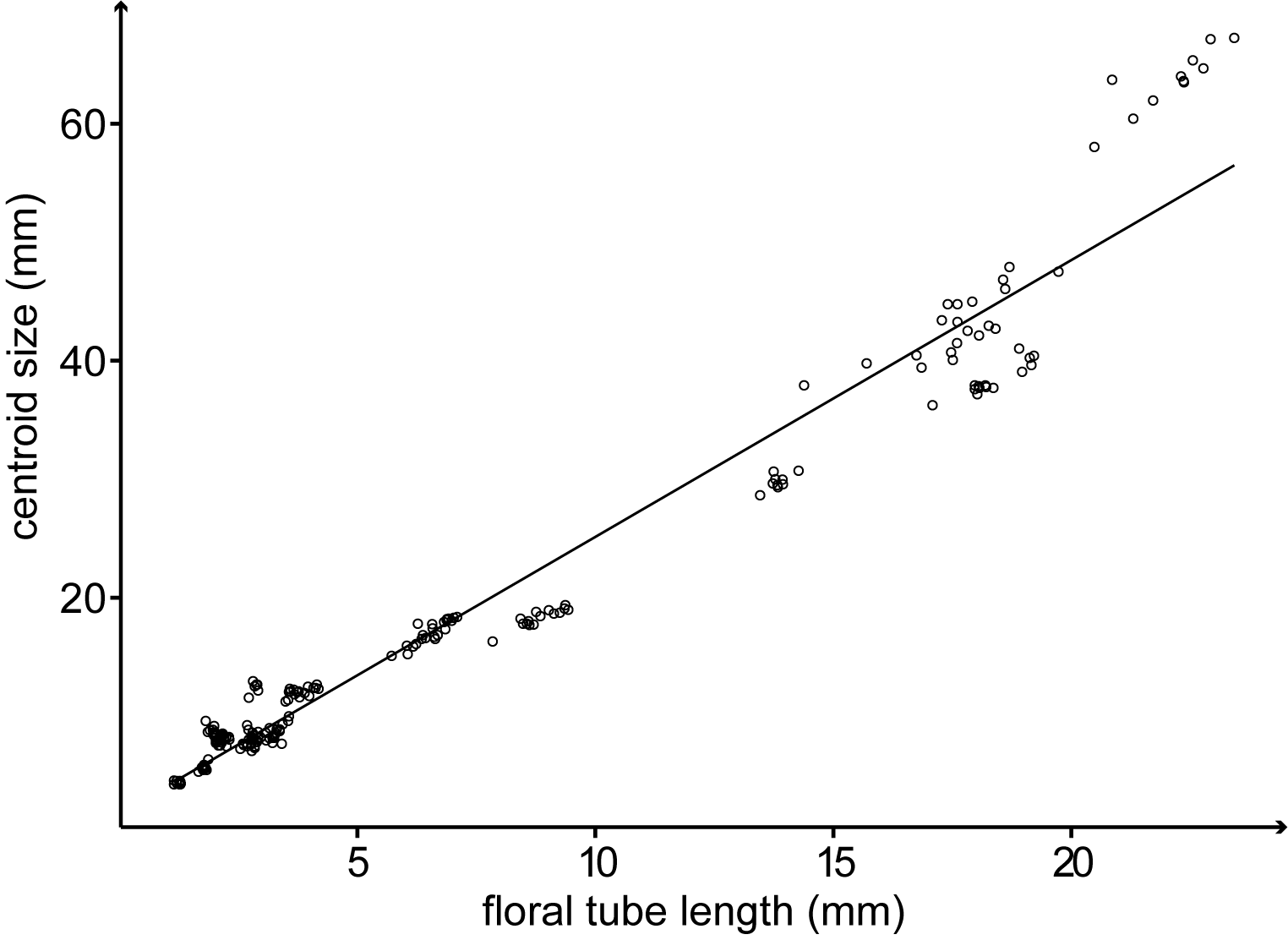

R2 = 0,96

P-value = 2.2 E-16
