## Supplementary figures and images for "Modularity and evolution of flower shape: the role of efficiency, development, and spandrels in *Erica*"

### Fig. S1

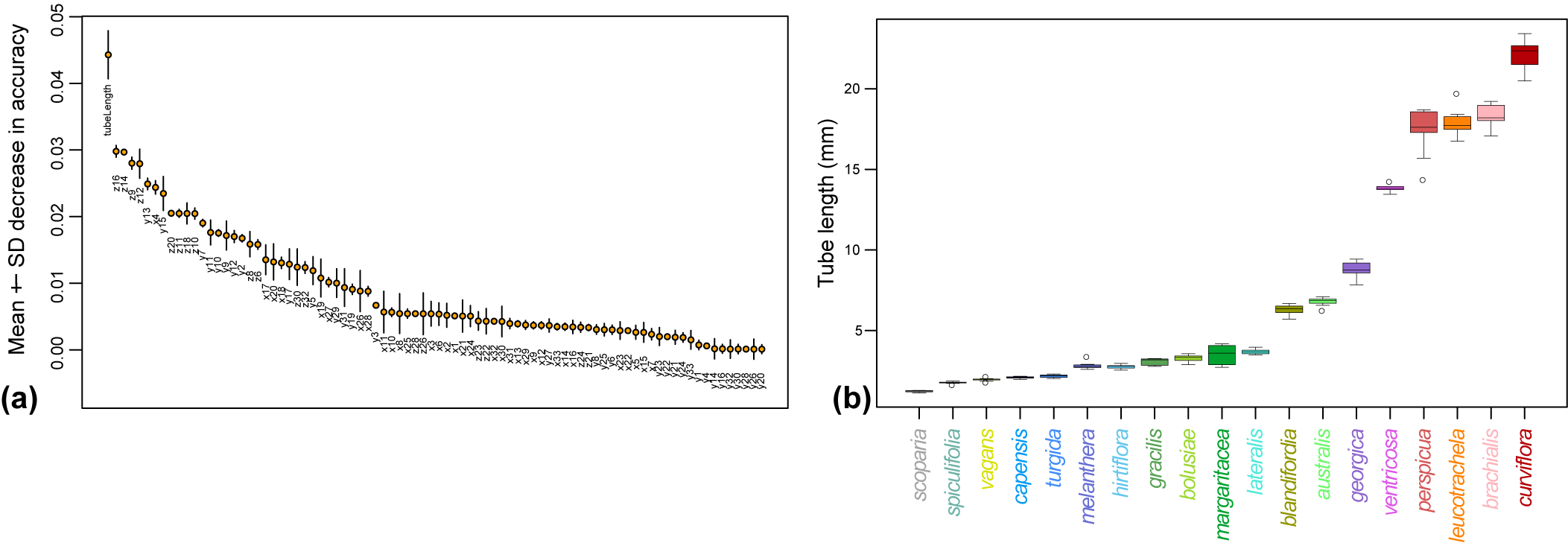

### Fig. S2

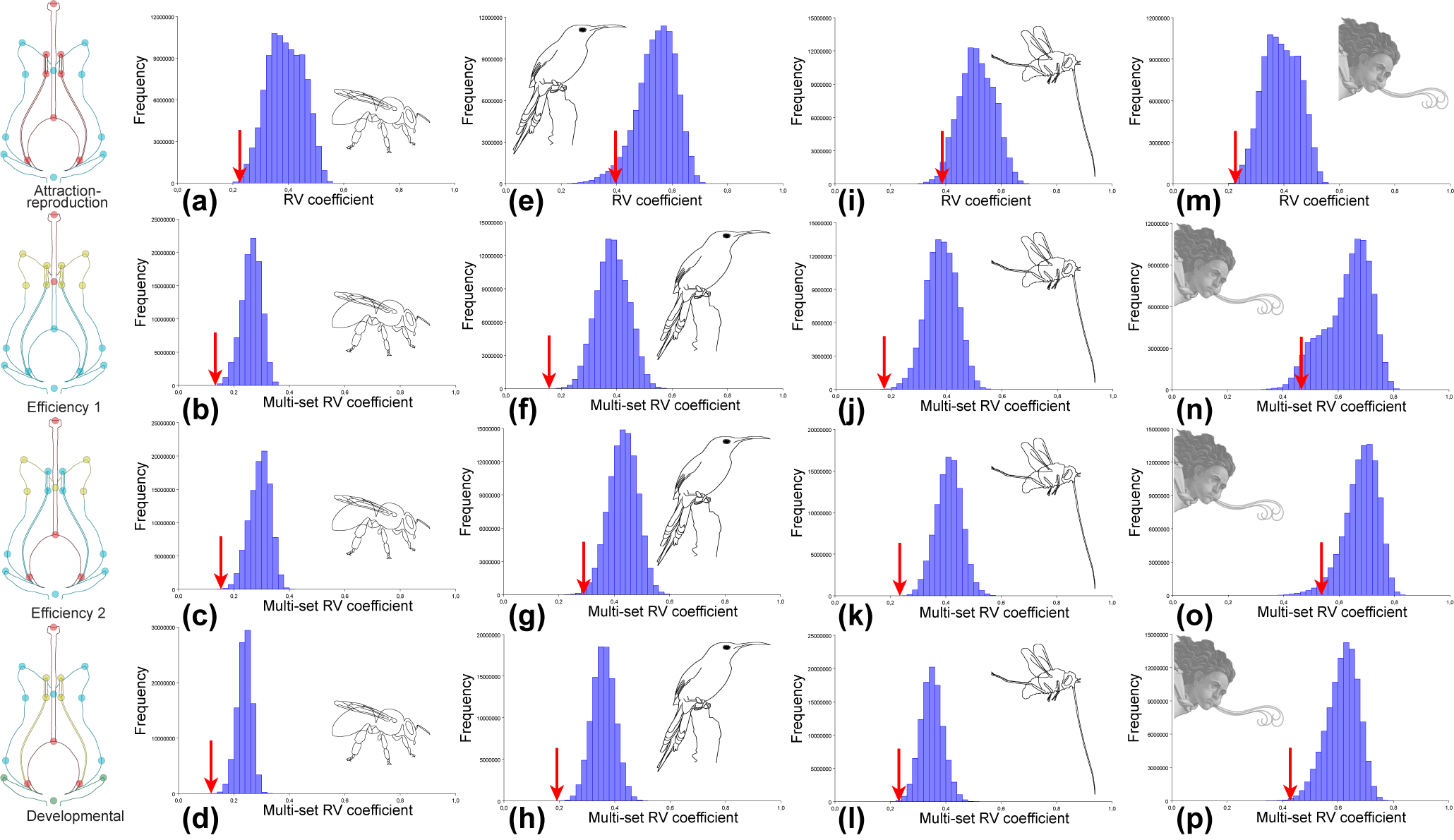

### Fig. S3

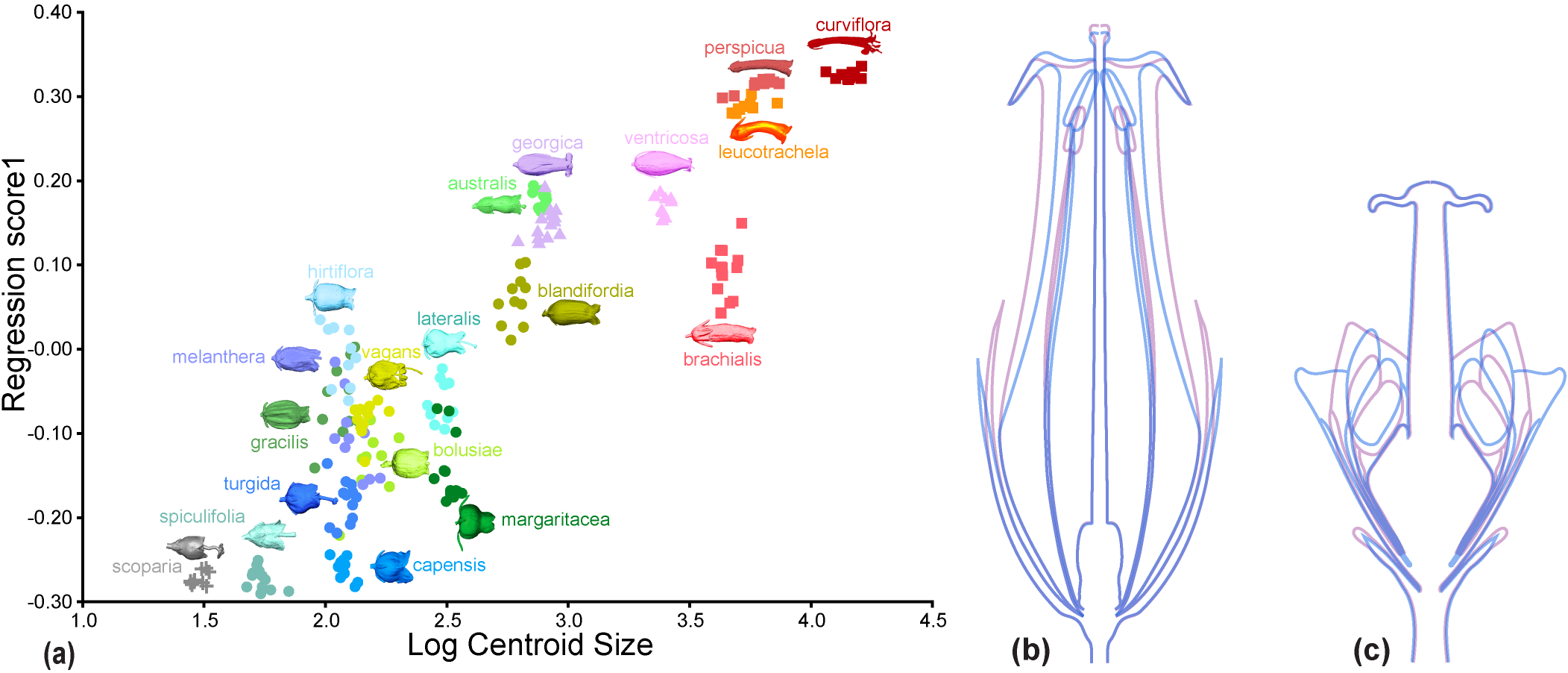
